# Accessing anti-HIV activity through the attenuation of USP18 activity: novel insights from molecular dynamic simulations, free-energy profiling, and multi-cellular inhibition assays

**DOI:** 10.1101/2024.06.23.600290

**Authors:** Lester T Sigauke, Jonathan Bvunzawabaya, Gabrielle Le Bury, Godwin A. Dziwornu, Saikat Boliar, David W Gludish, David G Russell, Krishna Govender, Grace Mugumbate, Nyaradzo Chigorimbo-Murefu

**Affiliations:** Division of Medical Virology, Department of Pathology, University of Cape Town, Rondebosch, South Africa; Department of Chemical Sciences, Faculty of Science and Technology, Midlands State University, Gweru, Zimbabwe; Microbiology and Immunology, College of Veterinary Medicine, Cornell University, Ithaca, New York, United States of America; Department of Chemistry, University of Cape Town, Rondebosch, South Africa; Department of Chemical Sciences, University of Johannesburg, Doornfontein Campus, Johannesburg, South Africa; International Centre for Genetic Engineering and Biotechnology, Cape Town, South Africa

## Abstract

The feasibility of achieving anti-HIV activity from the attenuation of USP18 activity was explored for the first time. A cheminformatic survey demonstrated that the current known USP18 isopeptidase inhibitors are derivatives of a bis-aryl pyranone scaffold that possesses undesirable toxicity profiles. Molecular modelling approaches applied to these active bis-aryl pyranones isolated the likely mechanism that perturbs the isopeptidase activity of USP18. Molecular dynamic simulations and free-energy profiling showed that induced-fit effects on the catalytic triad and the IBB-1 domain residues of USP18 drive a reversible non-competitive isopeptidase inhibition mechanism. Proof-of-concept multi-cellular HIV inhibition assays demonstrate the utility of achieving anti-HIV-1 activity from attenuating the activity of USP18 using small molecules. This study motivates for the pursuit of scaffolds that target the allosteric site of USP18, fine-tuning the IFN response as a strategy to enhance the natural control mechanisms that lead to an antiviral state potentially curing viral infection.

## Introduction

Although certain interferon stimulated genes (ISG) exist for the express purpose of dampening inducible interferon (IFN) responses as a natural control mechanism to prevent systemic inflammation, HIV-1 also attenuates this IFN signal cascade rendering it insufficient to resolve infection (1–4). The role of IFN signaling in HIV infection is thus complex and interventions that target the IFN response regulating ISGs remain underexplored. It has been speculated that fine-tuning of the IFN induced restriction factors that counter viral replication could potentially provide a pathway to the long-term control of HIV if their upregulation could be achieved in a limited focused way such as is demonstrated by IFNα2 (1). Achieving this effect through sustained expression of IFN however has thus far not yielded positive results in chronic HIV patients despite the lack of detrimental effects associated with the risk of increased systemic immune (5,6). As interferon therapy studies in chronic Hepatitis B virus have demonstrated distinct antiviral activities from different IFNα subtypes it is thought that perturbing the IFN pathway in HIV patients could lead to a similar antiviral state without the detrimental effects of sustained IFN expression (7).

Computer-aided drug discovery approaches such as molecular dynamics have contributed to the rapid search of small molecule candidates as well as the isolation of likely mechanisms that inhibitors use to perturb known targets (8–10). We aim to perturb the activity of Ubiquitin-specific peptidase 18 (USP18) because it has been identified as a dominant ISG that regulates the IFN response (11). USP18 is a 43 kDa cysteine isopeptidase that serves a complicated role of reversing the ISGylation of viral proteins as well as a non-catalytic function that regulates the inflammatory response (12–14). The non-catalytic function is due to USP18 interactions with interferon α/β receptor 2 (IFNAR2) which leads to the attenuation of IFN signaling preventing the detrimental impacts that arise from elevated and sustained inflammation (15–17). The catalytic USP18 role is related to a reversal of the post-translational modification of proteins by ISG15 (ISGylation). ISGylation regulates the activity of target proteins as part of the type I IFN mediated response that initiates the ubiquitin-proteasome system to trigger the anti-viral response (12,18,19). The catalytic role of USP18 reverses this ISGylation and affects the IFN response by dampening the anti-viral response (11). The inter-play of the ubiquitin-proteasome system is observed in SARS-CoV, where viral proteins mimic and suppress ubiquitin while other viral proteases have been shown to reverse the ISGylation of viral proteins reducing viral clearance (13,19).

Inhibiting the expression of USP18 has been shown to impact the survival of HIV-1 infected CD4 T cells, while an anti-viral IFN effect in liver cells infected with Hepatitis B and C resulted in viral clearance (20–22). This multifaceted function of USP18 as a deISGylating enzyme and a modulator of the IFN response has led us to hypothesize that approaches that selectively target either the catalytic isopeptidase or the non-catalytic activity of USP18 will lead to therapeutic strategies that can be applied in various human diseases such as HIV. Bis-aryl pyranone scaffolds have been shown to have nonselective and unclassified USP18 inhibitory potential in antineoplastic applications (23). Cersosimo et al (2015) synthesized two series of a previously identified non-selective isopeptidase inhibitor (N-SII) **G5**. The first series of the 12 compounds are based on the bis(arylidene)tetrahydrothiapyran-4-one 1,1-dioxide scaffold of **G5** with variations in the substituents on the aromatic rings. The second series are eight nitrobenzylidene pyranones acquired from a substitution of the **G5** sulfone group by different groups leaving the 4-nitro substituents on the aromatic rings fixed. From these synthesized **G5** analogues, only **2c** and the **G5** were tested in their cell based USP18 assays against the isopeptidase activity. The absence of the accumulation of free cleaved GFP led them to suggest that the observed USP18 inhibition was due to USP18’s reduced isopeptidase activity on the GFP tagged ISG15 chimera substrate. Both **2c** and **G5** were shown to inhibit the USP18 deISGylation of ISG15 tagged substrates with IC_50_ of 21.20 and 33.23 μM, respectively.

We report the first attempt at acquiring anti-HIV activity from agonists tuned to dampen the IFN response through the perturbation of specific ISG factors such as USP18. Anti-HIV therapeutics that target ISGs such as USP18 have the potential to serve as a natural control mechanism for inflammation and cell activation by enhancing end stage IFN signalling. To date this is the first study that has attempted to pursue this approach to anti-HIV activity.

## Materials and methods

### Synthesis of compounds

Compounds were synthesized as previously described and the chemical reagents were procured from Combi-Blocks Inc (23). All solvents were anhydrous or of analytical grade.

### Cheminformatics of the pyranone series

Absorption, distribution, metabolism, and excretion (ADME) properties of 13 pyranones were profiled using SwissADME server (24,25). The evaluation included parameters such as physicochemical properties, lipophilicity, water-solubility, pharmacokinetics, drug-likeness, and medicinal chemistry properties. The Brain Or IntestinaL EstimateD permeation method (BOILED-Egg) from SwissADME was implemented to graphically show the passive gastrointestinal absorption (GI) and blood-brain barrier penetrant (BBB) properties of the compounds respectively (25). ADMETlab 2.0 was used to compute toxicity properties such as hERG blockage, Human hepatotoxicity (H-HT), drug induced liver injury (DILI), Ames toxicity, rat oral acute toxicity (ROA) and carcinogenicity of the pyranones (26). Metabolic transformation products predictions were done using BioTransformer 3.0 (27).

### Metalated USP18 model building

In the absence of a crystal structure of the human USP18 protein, a human model of the USP18 protein was obtained from the SWISS model database (24). Model Q9UMW8 which had a 74.4 % sequence similarity to the mouse 5CHV crystal structure model was downloaded for this study. This USP18 homology model structure was submitted to the Metal-Ion Binding Site server to estimate the binding site region for the Zn^2+^ ions and the pose of the metal ion to acquire the metalated USP18 (28,29). new

### Molecular docking

The surface of the USP18 model was treated with the SiteMap tool excluding shallow solvent exposed pockets from its search to identify pockets that can be perturbed by small molecules (30). The SiteMap algorithm ranked the identified sites according to a SiteScore scoring function. The degree of discrimination of each site was determined by setting up grids for virtual screening of all the ligands at the centre of each of the sites. The 13 pyranone small molecule derivatives were constructed using the Marvin Sketch web tool and the ligands were prepared by the LigPrep workflow using the following parameters (hydrogens added and pH of 7.0 and a maximum of 32 stereoisomers for each ligand) (31,32). The docking methodology relied on the use of XP Glide docking followed by a Quantum Polarized ligand docking (QPLD) workflow (33–35). The QPLD docking workflow made use of the Jaguar Accurate method for the calculation of the ligand charges of each pose during the redocking stage. The QPLD output was limited to a maximum of 5 poses for each ligand from a generous RMSD threshold of 2Å to separate the different conformers. The total number of poses for each docked ligand using this threshold to evaluate the number of modes indicated the degree of discrimination of each site for each ligand. The 5.2 million ZINC15 compounds that satisfied the following criteria: MW greater than 325 g/mol and less than 425 g/mol, possessing 3D representations obtained at pH 7.4, anodyne reactivity to eliminate PAINS assay patterns, and purchasable compounds that are charged between +2 and -2. These ligands/compounds were downloaded in tranches from the online server using download scripts (36,37). These ligands were prepared using LigPrep before Glide XP docking with a grid box centered on the favored allosteric pocket of site 2. The ligands were ranked according to their Glide score and the top 200 candidates were selected for filtering based on the ZINC15 vendor databases to identify purchasable compounds. An in-house scoring metric ranked the purchasable compounds based on 29 features acquired from the SwissADME portal describing their solubility, pharmacokinetics, drug-likeness, and medicinal chemistry properties to prioritize compounds for purchasing.

### Single-trajectory molecular dynamics simulations

Prior to all molecular dynamics simulations Induced-Fit Docking (IFD) of the ligands in the metalated receptor with grid boxes centred on the allosteric pocket of site 2 was used to prepare the starting conformations for the systems (38,39). The following steps were carried out during the IFD workflow: Protein Preparation (RMSD 0.18), Glide Docking (SP, 20 poses per ligand, OPLS2005), Determine Residues for Refinement (Distance cut-off of 5.0 Å), PRIME Active Site Optimization (OPLS2005), Scoring and Filtering (filter using Prime Energy, energy window of 30 kcal/mol, keep maximum of 20 poses per ligand), Glide Docking2 (XP, 1 pose per ligand, OPLS2005), and Scoring (Prime Energy). Complexes with the most optimal IFDScore were extracted and used as the starting configuration for the single trajectory simulations for each system. Desmond single trajectory molecular dynamics simulations under constant pressure were performed to interrogate the protein-ligand interactions of the systems in solution (40). The systems were prepared in an orthorhombic box of buffer size 10 Å, solvated by TIP4P waters, and neutralized by adding sodium and chloride ions when necessary. The OPLS2005 forcefield was used for system preparation and production (41). Prior to the start of the 100 ns of NPT unrestrained production simulations, 160 ps of a standard NPT relaxation protocol was used to get each system to a temperature of 300 K and an isotropic pressure of 1.013 bar.

### Multi-trajectory molecular dynamics simulations

A multi-trajectory approach to interrogating the free-energy profile of the complexes made use of biased pulling simulations that were sent through to umbrella sampling of coordinates that trace the pulling reaction coordinate (42). These simulations were performed in GROMACS (version 2018.8) and made use of a CHARMM36 (version July 2021) forcefield for the atom parameters (43,44). Molecular topologies were acquired from single trajectory protein-ligand systems. The ligand topologies were acquired from the automated CHARMM General Force Field program (version 1.0.0) using valence-based bond orders (45–47). Stream files of the CHARMM36 protein topologies and the CGenFF ligand topologies were converted to GROMACS format and merged manually before the system was solvated using TIP4P waters, and neutralized using chloride ions (46). The systems were minimized and equilibrated before steered/biased NVT production simulations were propagated. The steered pulling simulations were performed on a box that was extended along the z axis, allowing the ligand to be pulled away from the receptor at a rate of 0.01 nm ps^-1^ with a force of 1000 kJ mol^-1^ nm^-2^ until greater than 5 nm separation was achieved. Windows separated by center of mass (COM) distances between the ligand and protein of either 0.2 nm or 0.12 nm defined the configurations that sample the reaction coordinate from the biased unbinding trajectories. For each system about 20 configuration files were selected from these windows as the steps along the bind reaction path. These files/steps were the starting points for the multi-trajectory simulations with equilibration being performed before NVT production simulations of 1 ns at each step. For each system these simulations gave rise to an ensemble of overlapping configurations that traced the reaction path. A WHAM analysis applied over the entire ensemble of configurations for each of the systems gave the potential of mean force (PMF) that models the unbinding path (48). For each ensemble, these PMF curves gave access to the free-energy of bind (ΔG) of the binding/unbinding path from the extremes of the reaction path traced by the trajectory. The slope of this PMF curve and the COM windows gave the friction plots and compared the dissociation time in an analogous manner to the residence time.

### Multi-cellular HIV inhibition assays

#### Virus production

An HIV-1 infectious molecular clone, pWT/BaL (HXB3/BaL, Catalog 11414), was obtained from an NIH AIDS Reagent Program. The plasmid encoding HIV-1 molecular clone NL43-IRES-mCherry with R5-tropic env (BaL) was generated by replacing the EGFP sequence with that of Gaussia luciferase (gLuc) in the original plasmid pBR43IeG-nef+ R5env (a kind gift from Thorsten R. Mempel, Massachusetts General Hospital, Boston). Replication-competent VSV-G pseudotyped HIV-1 virus was prepared by co-transfecting 293FT cells with the respective HIV-1 molecular clone and VSV-G expression plasmid using Lipofectamine 3000 reagent (Life Technologies). Transfection media was replaced with fresh antibiotic free DMEM media 8 h post-transfection. After 72 h, cell culture supernatant containing HIV-1 virus was harvested, centrifuged to remove cell debris, passed through 0.45-µm filter, and stored at −80 °C in aliquots. Multiplicity of infection 0.05 of virus was used to infect human MDMs.

### Human monocyte derived macrophage differentiation

Human monocytes were obtained from peripheral blood mononuclear cells of healthy individuals by counter current centrifugal elutriation with an average purity of >97% (Elutriation Core Facility, University of Nebraska Medical Center). Monocytes were maintained in DMEM media supplemented with 10% human serum, 100 U/mL penicillin, and 100 µg/mL streptomycin (Invitrogen). Cells were cultured at 37 °C with 6% CO2 for 7 – 8 days to fully differentiate into macrophages (MDMs).

## Results and discussion

The goal of this study was to demonstrate whether the anti-USP18 inhibition strategy could fine-tune the IFN response by enhancing natural control mechanisms that lead to viral clearance. Illuminating the structural dynamics of USP18 inhibition by the known actives will assist in the development of new anti-HIV therapeutics with greater potency.

### Presenting the USP18 active bis-aryl pyranones 2c and G5

Although the activity of compounds **2c** and **G5** are known the mechanisms by which they perturb USP18 have not been determined. In the absence of a crystal structure of the human USP18 protein in complex with an active USP18 inhibitor it was pertinent to first predict the likely binding site of the active anti-USP18 pyranones **G5** and **2c**. This would enable elucidation of the mechanism employed by these antagonists to inhibit USP18. Molecular modelling approaches coupled with an interrogation of the free-energy profile of binding at this site were used to guide the selection of this site and propose a mechanism based on the known activities of these pyranones.

### Isolating the binding site of the USP18 active bis-aryl pyranones using docking

The SiteMap workflow identified and ranked according to their SiteScore 5 pockets on USP18 that are accessible to small molecule inhibitors (Fig 1).

**Fig 1:**
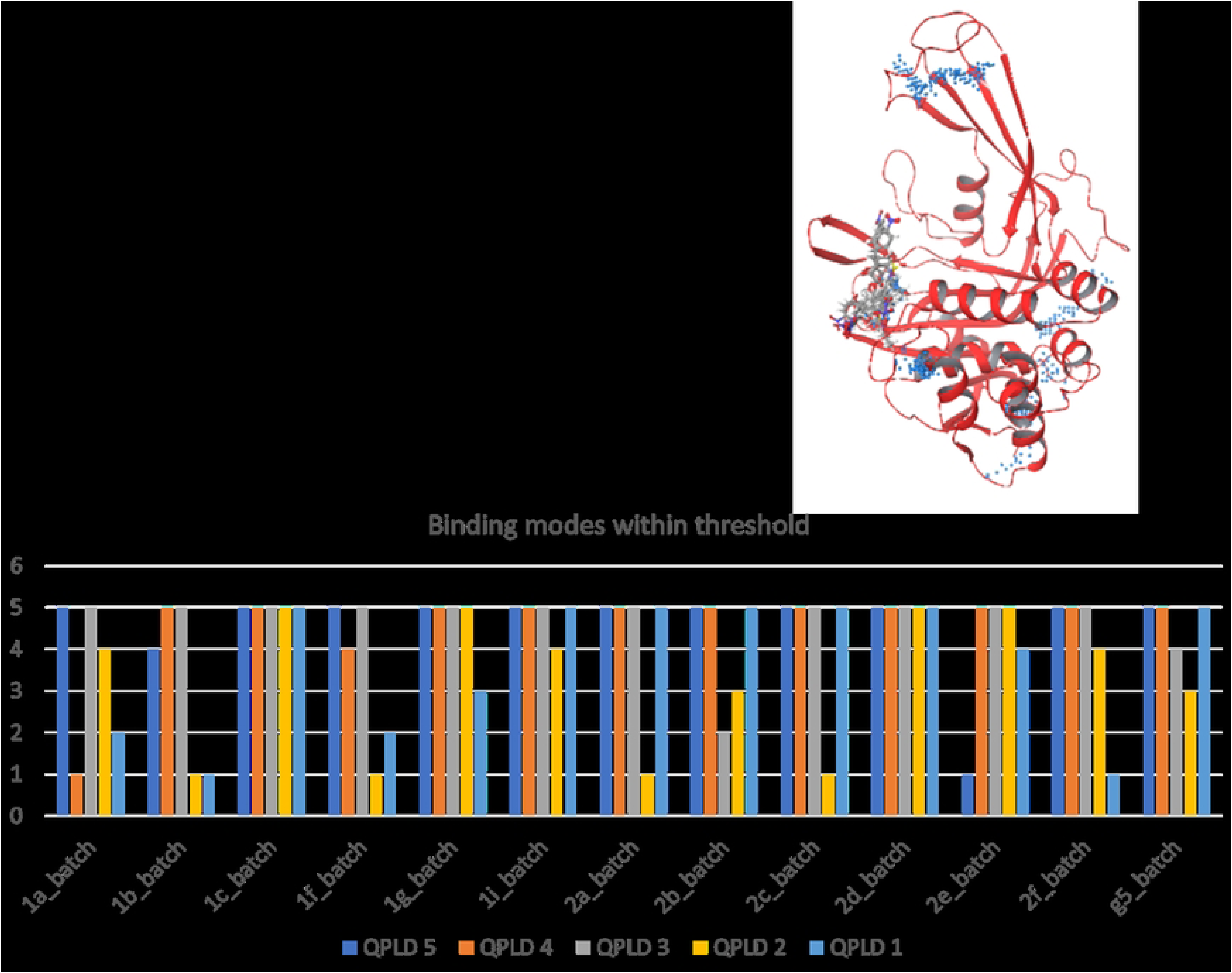
Docking on USP18. A. Ranked docking sites on USP18 identified by SiteMap. B. Ribbon structure of USP18 with map of small molecule docking sites (pseudo atoms are blue spheres). Small molecules shown are docked in site 2. C. Binding modes from poses of QPLD docking that satisfy the RMSD threshold for the 13 pyranone ligands on each of the 5 USP18 sites.

Of the 5 sites identified by SiteMap, site 2 (1.029) was the pocket most optimal for small molecule access (Fig 1 A). Grids for QPLD of the 13 proposed pyranone ligands were placed at the centre of these 5 sites identified by SiteMap. The centre of site 2 overlapped with the centroid of residues **Ala299**, **Asp124**, **Gln122** and **Tyr348** of the human USP18 sequence from the Q9UMW8 homology model. The binding modes from QPLD of the 13 pyranone ligands showed that site 2 (QPLD 2) was the most discriminatory site. This site had discriminatory potential expressed as achieving less than 5 poses, the maximum allowed number of poses, for 9 ligands while site 1 discriminated for 6 of the 13 ligands (Fig 1 C). For the known USP18 actives site 2 biased **2c** to 1 binding mode and 3 modes for **G5.** Site 1 however had 5 or more modes for ligand **2c** and **G5**. The SiteMap rank and its ability to restrict the binding modes affirms the suspicion that site 2 is a binding pocket able to bias active ligands into active modes and is thus the binding pocket of these 2 active inhibitors. Structural studies such as saturation transfer difference NMR (STD-NMR) of the ligands in solution with USP18 would confirm these observations in the absence of solved crystal structures of the protein-ligand complexes.

### Using the free-energy profiles of the USP18 active bis-aryl pyranones to guide site selection

To understand how binding at site 2 impacts the functioning of USP18, complexes of USP18 docked with the two active pyranone ligands at this site were studied using molecular dynamics simulations. Based on the known activity of **2c** on USP18 compared to the activity of **G5**, there was a need to find a plausible explanation for the slightly improved activity of **2c** over **G5**. IFD of these 2 ligands at site 2 gave access to starting structures for Desmond simulations and a stabilized conformer from these simulations were used for multi-trajectory umbrella simulations using GROMACS to evaluate the free-energy profiles of binding/unbinding.

The ligand interaction networks from the Desmond simulations showed that throughout the 100 ns of production simulation **G5** directly made use of 7 intermediary water molecules in its binding at site 2 whereas ligand **2c** recruited only a single water molecule in its interaction (Fig 2 A, B). The reliance on fewer water molecules for the **2c** interaction reflected on its stable ligand conformation maintained during the simulation compared to that of **G5** (Fig 2 C, D). This observation is consistent with the QPLD binding mode observation that showed that **2c** bound at site 2 had only one binding mode whereas **G5** had 3.

**Fig 2:**
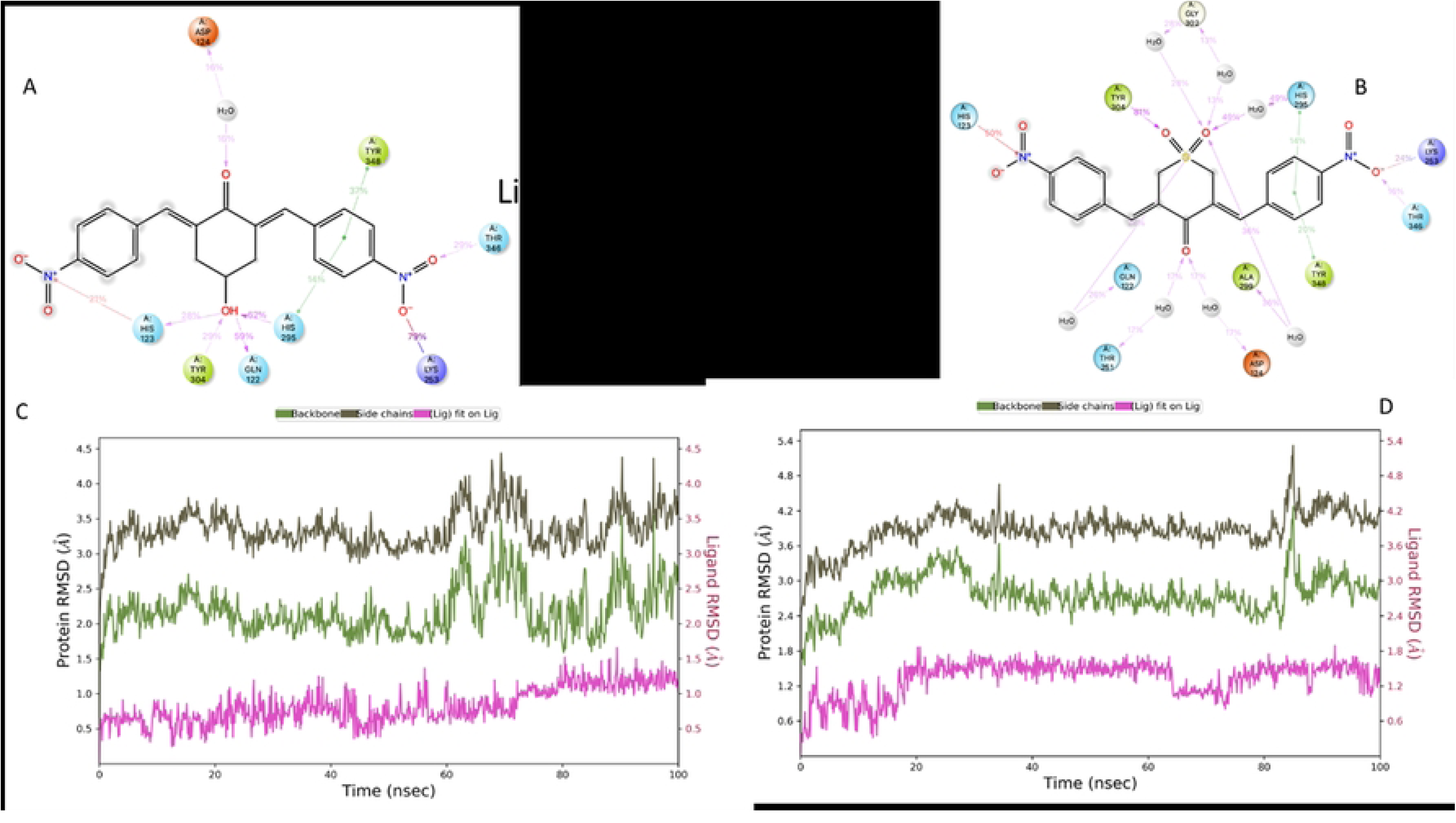
Ligand interaction network of ligand 2c (A) and ligand G5 (B) on site 2 during 100 ns of NPT Desmond simulations. RMSD plots of the ligand and the protein backbone and protein side chains of the USP18 complexes with ligand 2c (C) and ligand G5 (D) bound to site 2 during the 100ns of NPT Desmond simulations.

Interrogations of the free-energy profiles of the ligand complexes highlight the differences in the stabilities of the interactions of the 2 known USP18 actives at site 2 (Fig 4). Ligand **2c** (red line) dissociates from the active site later than ligand **G5** (blue) line (Fig 3 A, B). This difference in the dissociation is reflected by the PMF curves which estimated the free-energy of the binding reaction. Ligand **2c** (-10.55 kcal mol^-1^) has a slightly larger free-energy of binding than ligand **G5** (-10.08 kcal mol^-1^) (Fig 3 C, D). When taken together, the recruitment of intermediary waters from the ligand interaction network, the stability of the binding conformation from the RMSD, the dissociation time and the free-energy of binding, confirm that site 2 binding of **2c** and **G5** is responsible for the anti-USP18 activity observed by Cersosimo et al., 2015.

**Fig 3:**
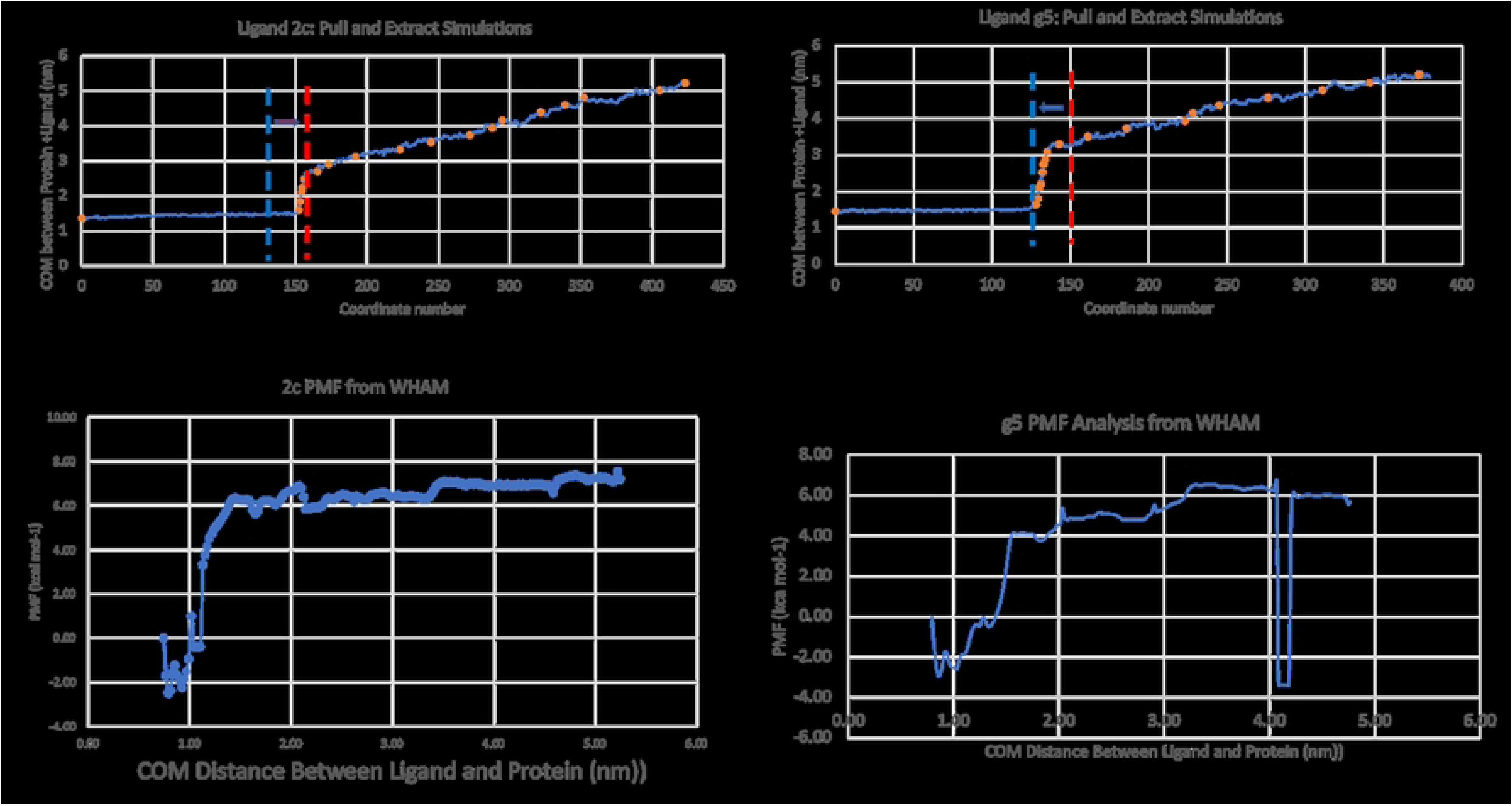
Free-energy profile plots. A., B. COM positions of trajectories from steered dynamics simulations of the ligand pulled away from USP18. Location of the asymmetric coordinate files that trace the ligand pulling trajectory for umbrella sampling of the dissociation path. Blue dash line shows the point of dissociation for ligand G5, Red shows the point of dissociation for ligand 2c. Arrows show the shift from the competing ligand. C., D. Potential of mean force (PMF) plots obtained from the WHAM analysis across the ensemble of configurations that trace the pulling trajectory of the dissociation path for ligand 2c and G5. ΔG_d_^2c^ = -2.50 + (-7.55) = -10.55 kcal mol^-1^ ; ΔG_d_^G5^ = -3.38 + (- 6.70) = -10.08 kcal mol^-1^ ; ΔG_d_^2c^ >= ΔG_d_^G5^.

**Fig 4:**
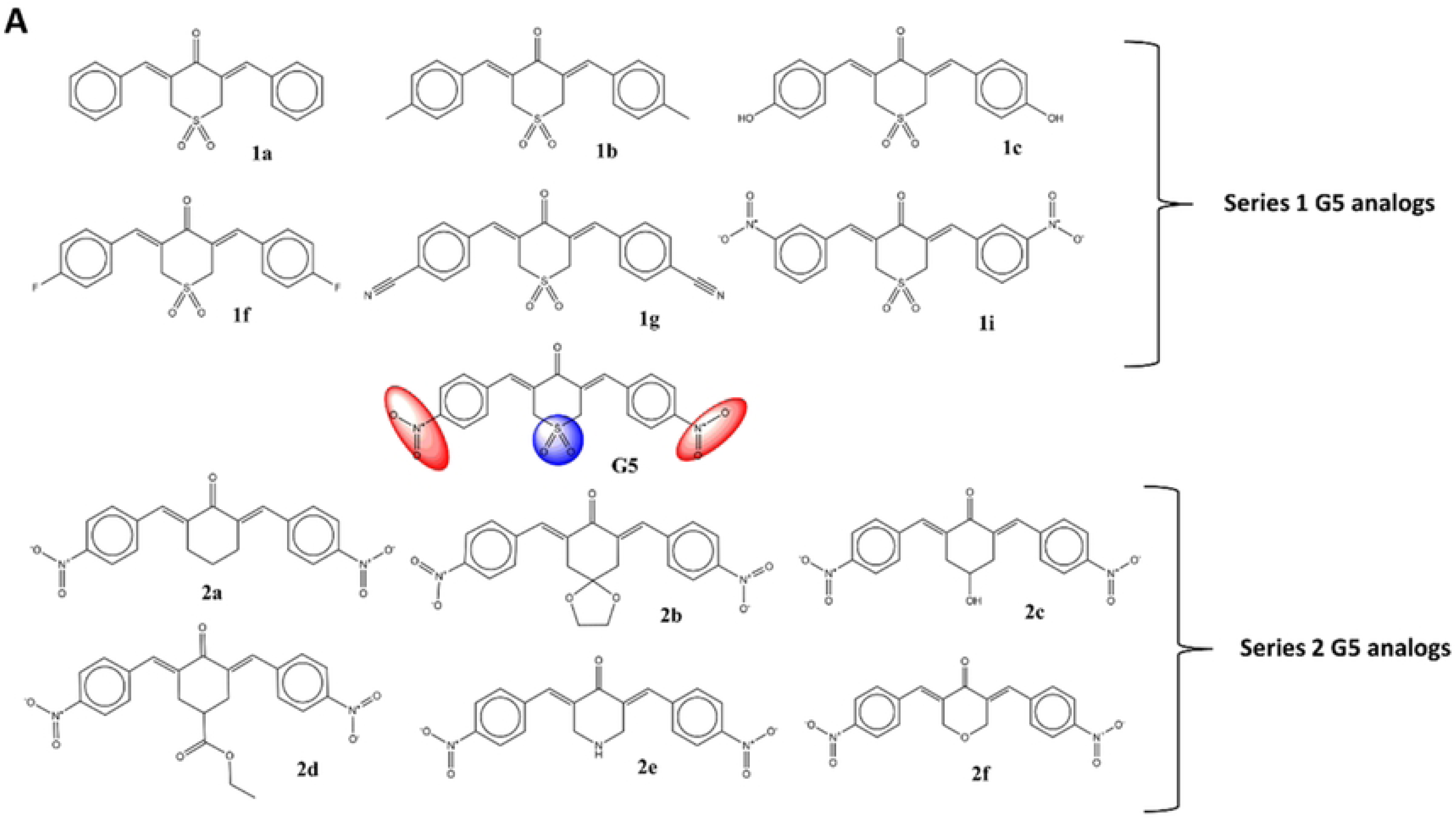
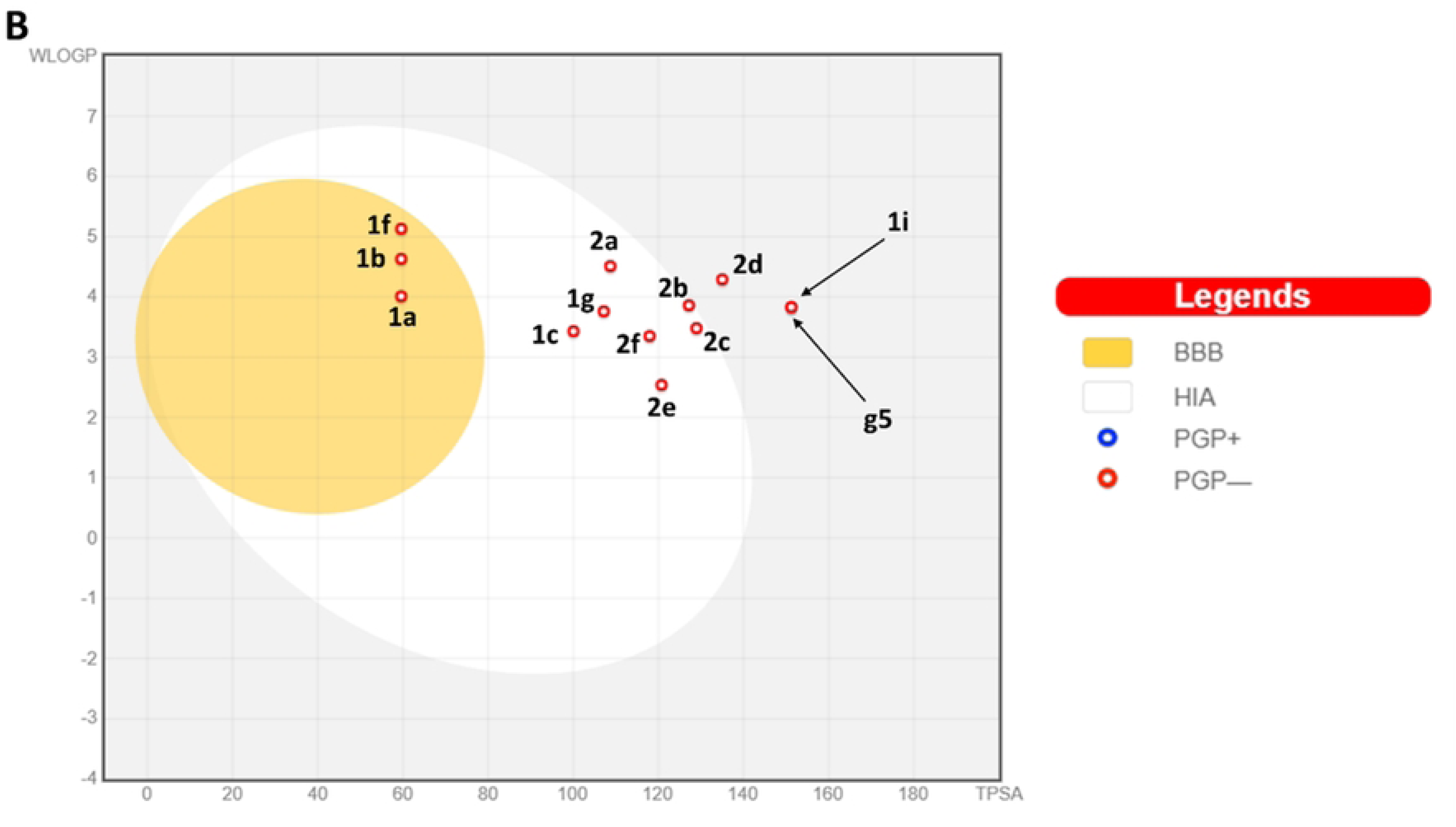
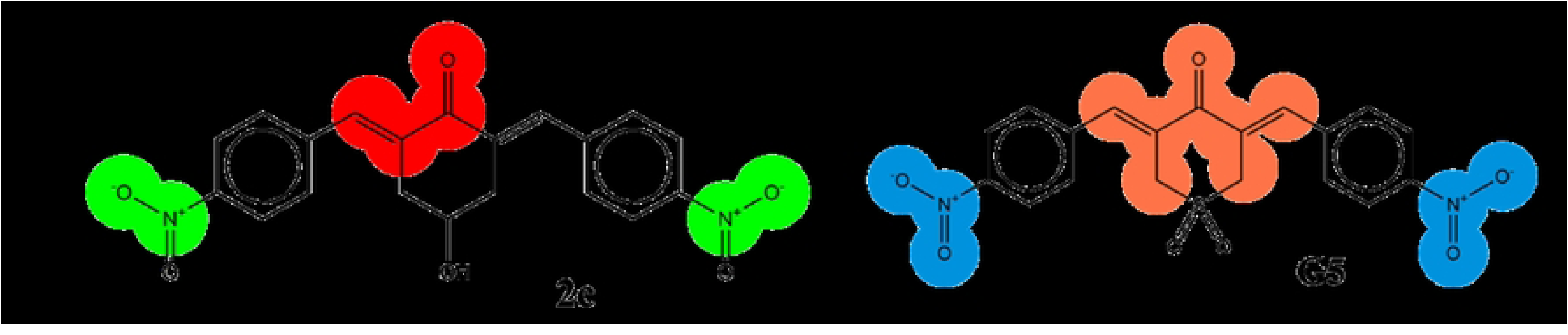
Cheminformatic analysis of pyranone derivatives. A. G5 analogues proposed by Cersosimo et al., 2015 used in this study. Series 1 analogues are based on variations in the substituents on the positions of the nitro group (red) of G5. Series 2 are based on the substitution of the sulfone group (blue) of G5. B. BOILED-egg model of the bis-aryl pyranones from the SwissADME server. The white region highlights compounds with high probability of GI absorption and the yellow highlights those with a high probability of BBB penetration. PGP-compounds are non substrates for P-gp. Compounds outside the grey region are not likely to be absorbed by GI and are not BBB permeant. C. PAINS and Brenk substructural alerts on 2c and G5. The ene_one_ene_A group (orange) is the only PAINS alert while the Brenk alerts present are the Michael acceptor 1(red), the oxygen_nitrogen single bond (green) and the nitro_group (blue).

### Presentation of the bis-aryl pyranone scaffold and its potential for pharmaceutical applications

It was pertinent to determine whether the bis-aryl pyranone scaffold was ideal for pharmaceutical applications. QSAR and machine learning derived models were used to predict the pharmacokinetic and toxicity profiles of the 12 Cersosimo derivatives of bis(arylidene)tetrahydrothiapyran-4-one 1,1-dioxide (bis-aryl pyranone), **G5**. All the bis-aryl pyranones that we examined possess two substituted aromatic benzene rings that are connected to a substituted cyclohexanone by double bond linkers. This connectivity is planar and reduces the fraction of sp^3^-hybridized carbons and the torsions within the small molecules reducing their flexibility (Fig 5 A). The 13 compounds have physicochemical properties that adhere to Lipinski and Ghose drug-like rules (Supplementary Table 1) (49,50). Although **1i**, **1d** and **G5** have TPSA values that disobey Veber, Egan and Muegge properties the rest of the pyranones have ideal bioavailability scores which would translate to good oral bioavailability (51–53). All our compounds have low and acceptable LogP values (**LogP _o/w_** < 5) with some being classified as soluble or moderately soluble (Supplementary Table 2).

**Fig 5:**
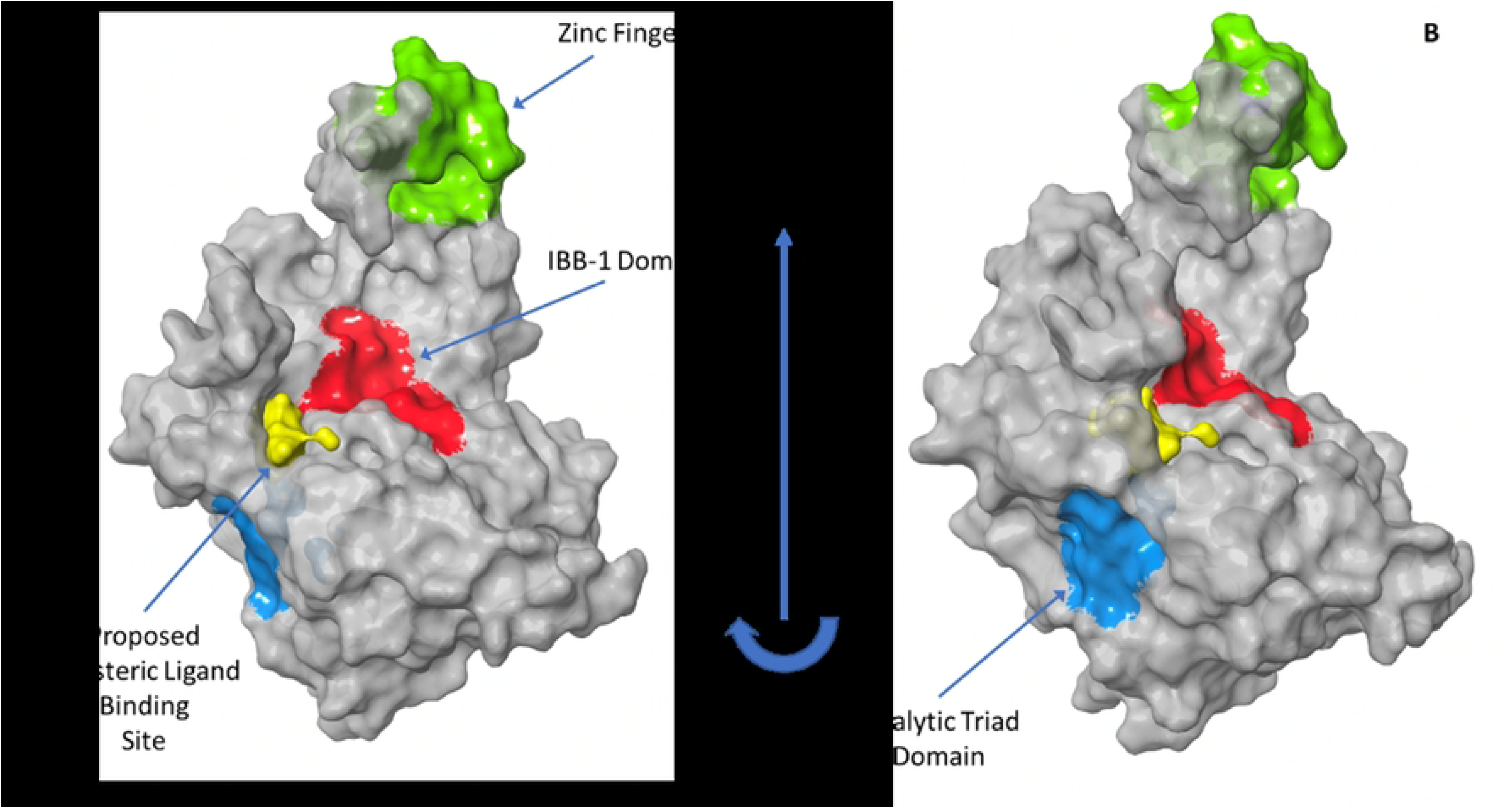
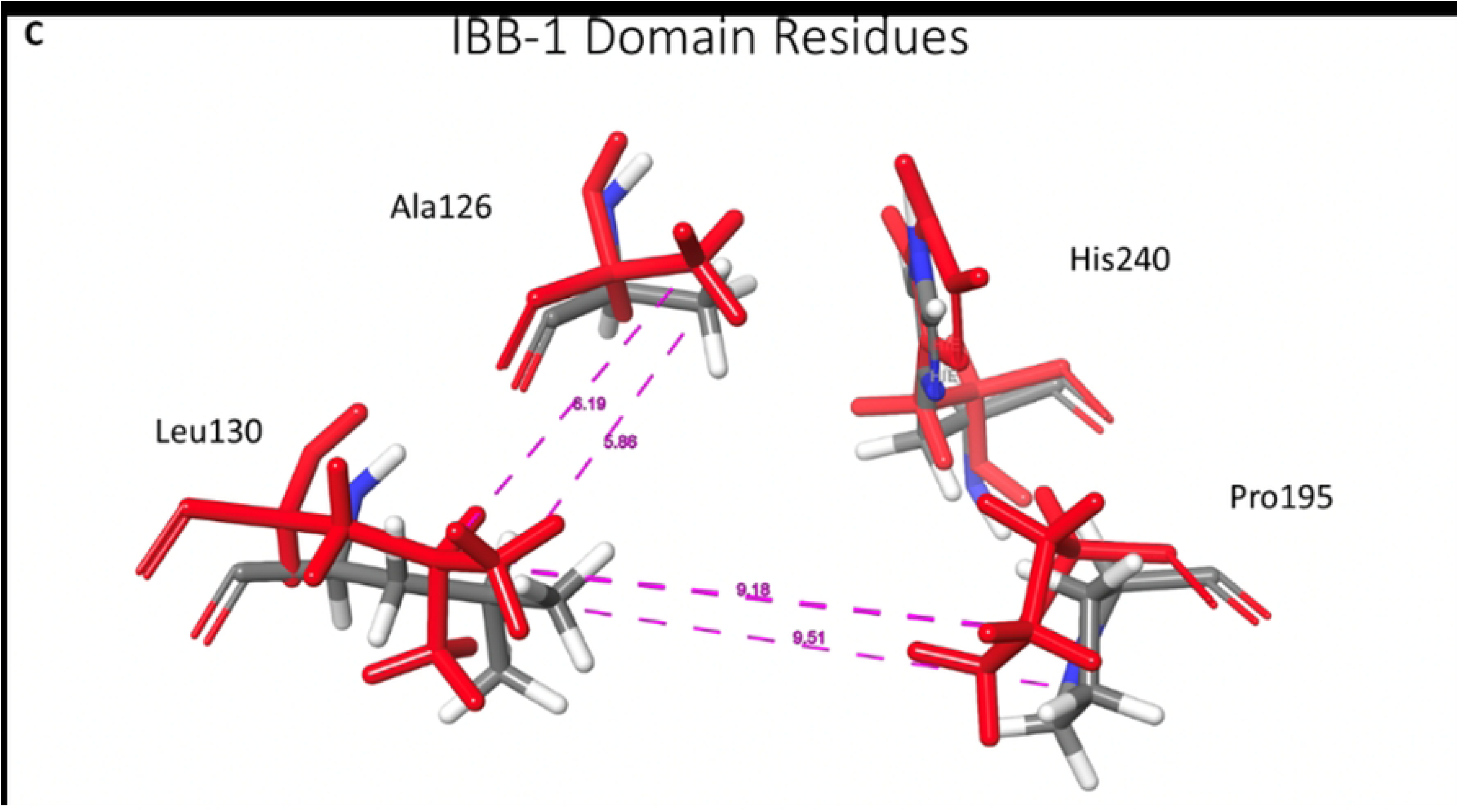
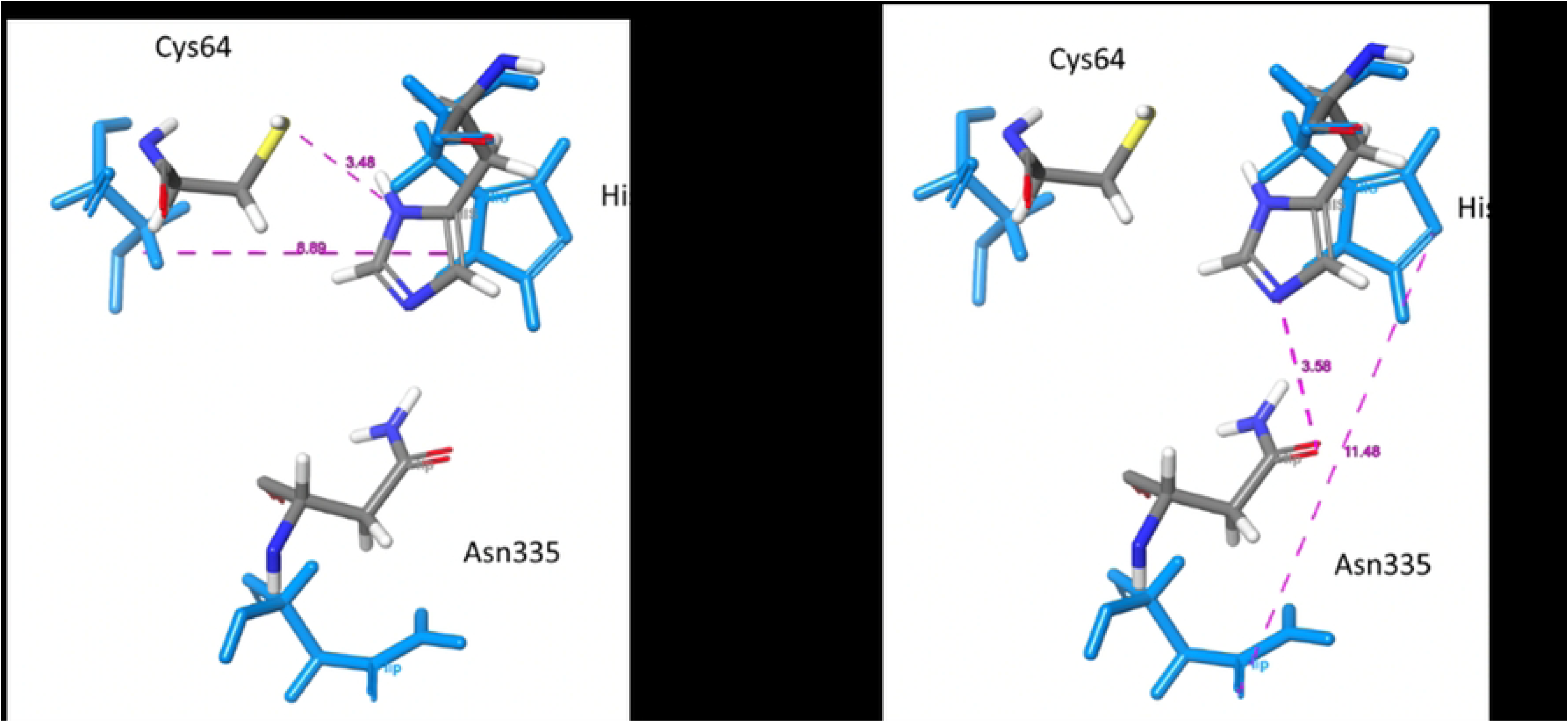
Perturbed USP18 domains. A, B Human USP18 surface. Green: Zinc finger domain, Red: IBB-1 domain, Blue: Catalytic triad domain, Yellow: location of Site2 pocket with surface around pseudo ligands from SiteMap. C. Overlap of IBB-1 domain residues in ligand bound with 2c conformation (Red) and ligand unbound conformation (cpk). D. Overlap of catalytic triad residues in ligand bound with 2c conformation (Blue) and ligand unbound conformation (cpk). Purple lines trace some of the bond lengths monitored from trajectories of the NPT simulations.

Apart from **1i**, **2d** and **G5**, the ADME profiles of these compounds predict high gastro-intestinal (GI) absorption (Fig 5 B). With exceptions to **1a**, **1b**, **1f** and **1g**, they have low polar surface areas (TPSA <90) and are not likely to penetrate through the blood brain barrier (BBB) (54) BBB permeant compounds such as **1a**, **1b**, **1f**, and **1g**, should be avoided as they carry the possibility of undesired pharmacological events associated with potential neurotoxicity (54) None of the pyranones are P-glycoprotein (P-gp) substrates, therefore their bioavailability cannot be affected by P-gp flux transporters (56). Unfortunately, all 13 compounds inhibit at least one of the Cytochrome P450 (CYP450) human isoforms CYP1A2, CYP2C19, CYP2C9, and CYP3A4 with none inhibiting the CYP2D6 isoform. Of this set **1c**, **1i**, **2b**, **2e** and **G5** have the best metabolic profiles as they inhibit only one CYP450 isoform. As CYP450 enzymes are responsible for metabolizing 80 % of drugs, therefore any compound that inhibits P450s is likely to be hepatotoxic (57). All 13 pyranones possess the **ene_one _ene _A** PAIN alert. Compounds **1a, 1b 1c, 1f** and **1g** contain the **michael_acceptor_1** as a Brenk alert while the nitro groups on **1i, G5** and **2a-2f** are flagged as additional Brenk alerts (Fig 5 C). These features are used to predict compounds that show false positive activity, metabolically instability, putative toxicity, chemically reactivity and poor pharmacokinetic properties (58,59). Although the **G5** analogs had low cardiotoxicity, they had a high probability of triggering drug-induced liver injury, while the nitro containing derivatives were flagged for mutagenicity and hepatotoxicity from the ADMETlab toxicity predictions (Supplementary Table 3)(60,61). Although several approved drugs in clinical use contain the nitro group this group is avoided due to its likelihood of triggering toxicity considerations (62). Overall, despite these compounds being druglike and orally bioavailable the pyranone scaffold makes them not ideal for medicinal and pharmaceutical applications. The metabolic and toxicity profiles of these compounds show that they possess CYP450 inhibition, carcinogenicity, and hepatotoxicity. Compound **G5** and **2c** possess significant structural alerts that would need removal if these scaffolds were to be pursued for further development. Undeterred by this negative outlook, this class of compounds however enables us to probe the role that antagonism of USP18 plays in the development of novel anti-HIV therapies due to the documented isopeptidase activity of **G5** and **2c**.

### Description of anti-USP18 activity which leads to anti-HIV-1 activity

Based on the observed anti-HIV activity (Fig 9) 8 compound systems were selected to guide the development of a hypothesis that explains the role that USP18 activity could have on anti-HIV activity. The ligand systems selected were ligand **1a**, **2a**, **2c**, **2d**, **G5**, **Chem46** (acquired from ChemDiv), **Mol33** and **Mol71** (acquired from Molport).

### Allosteric inhibition

The USP18 active ligands (**2c** and **G5**) are hypothesized to have an allosteric perturbation of USP18 owing to the proximity of their preferred binding pocket, site 2, to the residues that affect USP18 activity (Fig 6). It is possible that the allosteric impact could arise from changes within the IBB-1 domain or changes within the catalytic triad region brought on by binding of these active small molecules at the proposed allosteric site 2. If the activity is due to impacts on the IBB-1 domain, these changes will need to be significant enough to successfully impede the binding of USP18 to the ISGylated target substrate prior to proteolysis. Where the activity is due to changes to the triad residues, such perturbations would need to stall or reduce the proteolysis reaction that cleaves the substrate after the ISGylated target binds successfully to the inhibitor bound USP18. An additional allosteric non-enzymatic impact could result in limiting the upstream binding of USP18 to INFAR2 and STAT2 that serves to dampen the interferon cascade (Fig 1). It is, however, not immediately apparent how limiting this interaction would result in the observed viral clearance. Current understanding, instead, is that binding of USP18 to IFNAR2 reduces IFN signaling and dampens expression of antiviral ISGs such as USP18 and ISG15 resulting in a decrease in the rate of ISGylation and thus viral clearance. Isolating the likely mechanism employed by these ligands will assist in the development of inhibitors that enhance the perturbations that are leading to viral clearance.

**Fig 6:**
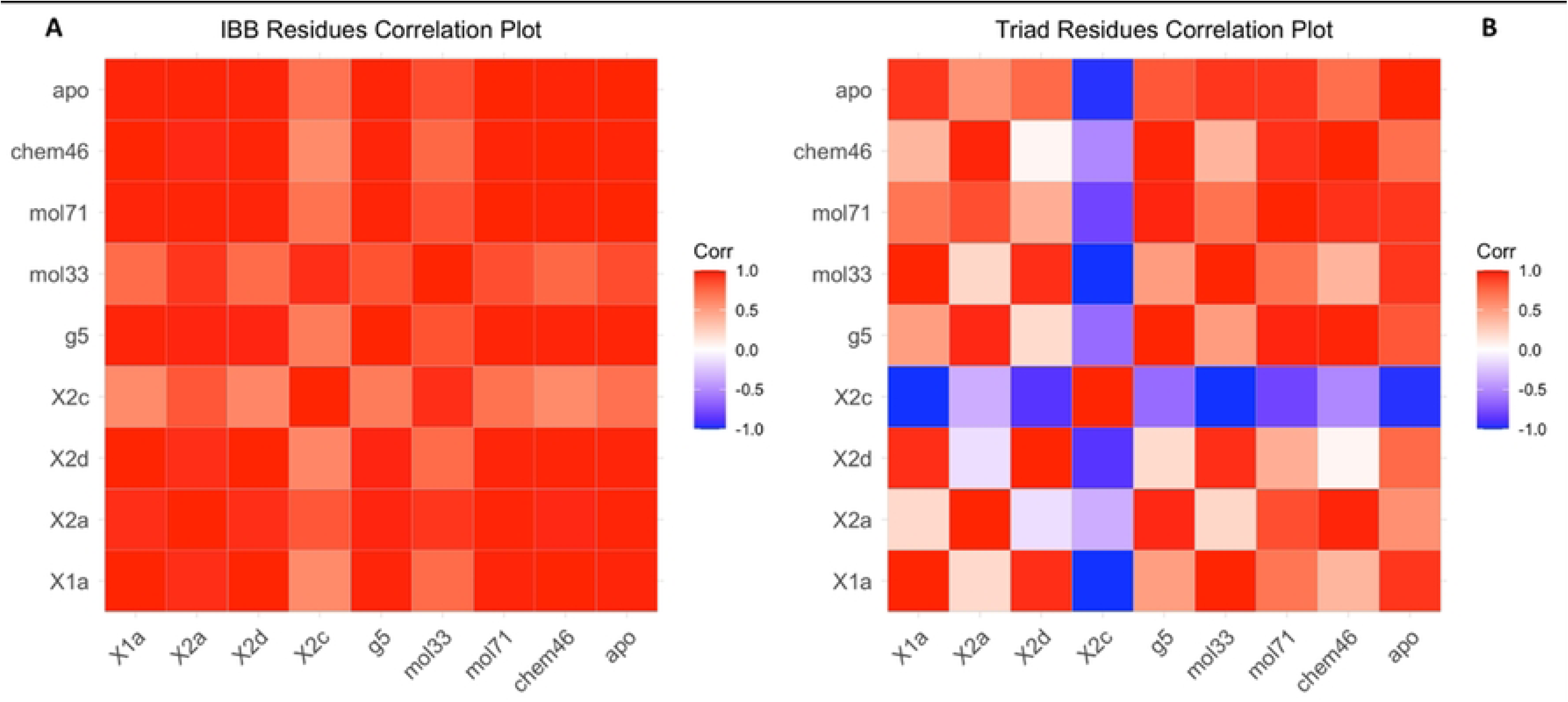
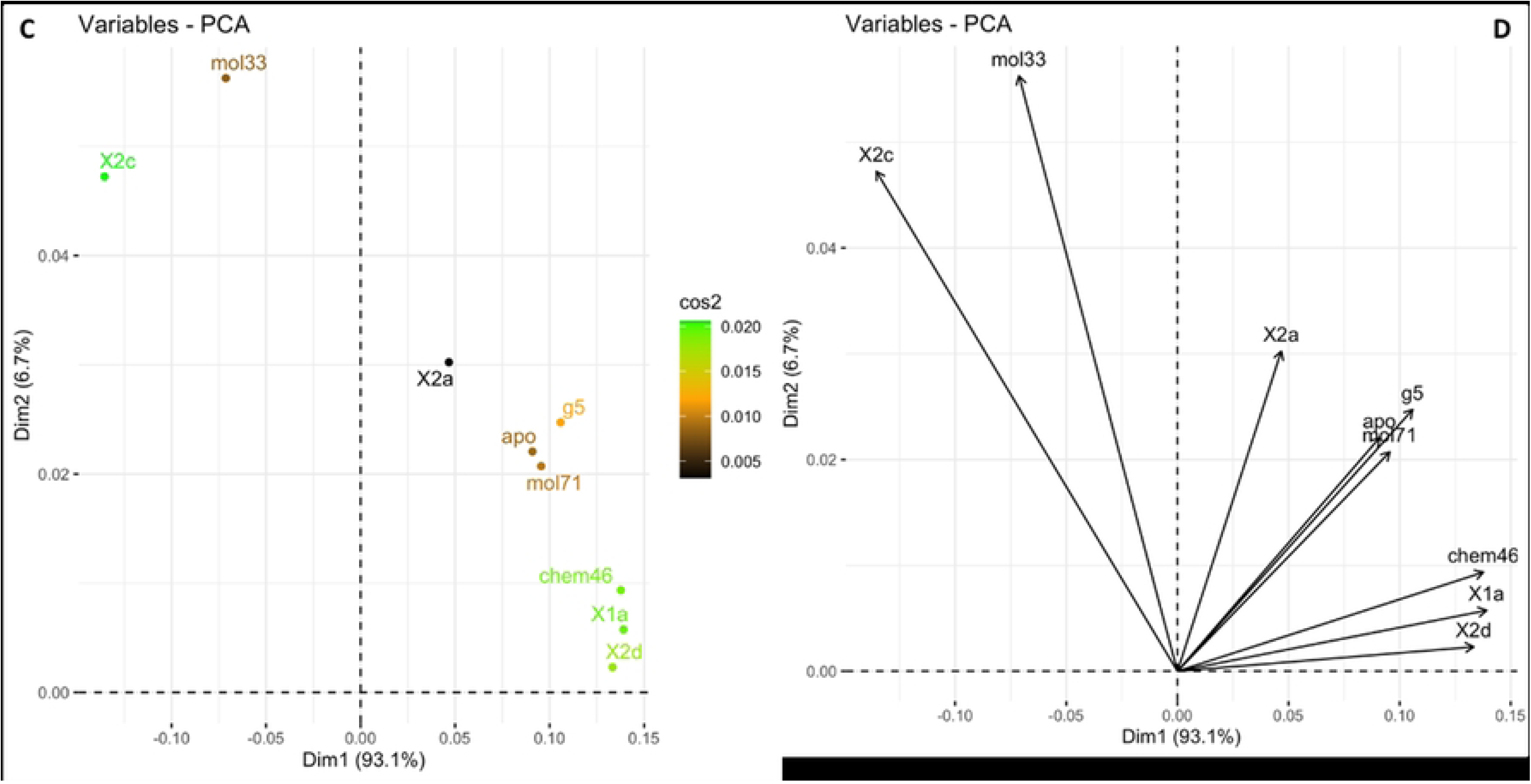
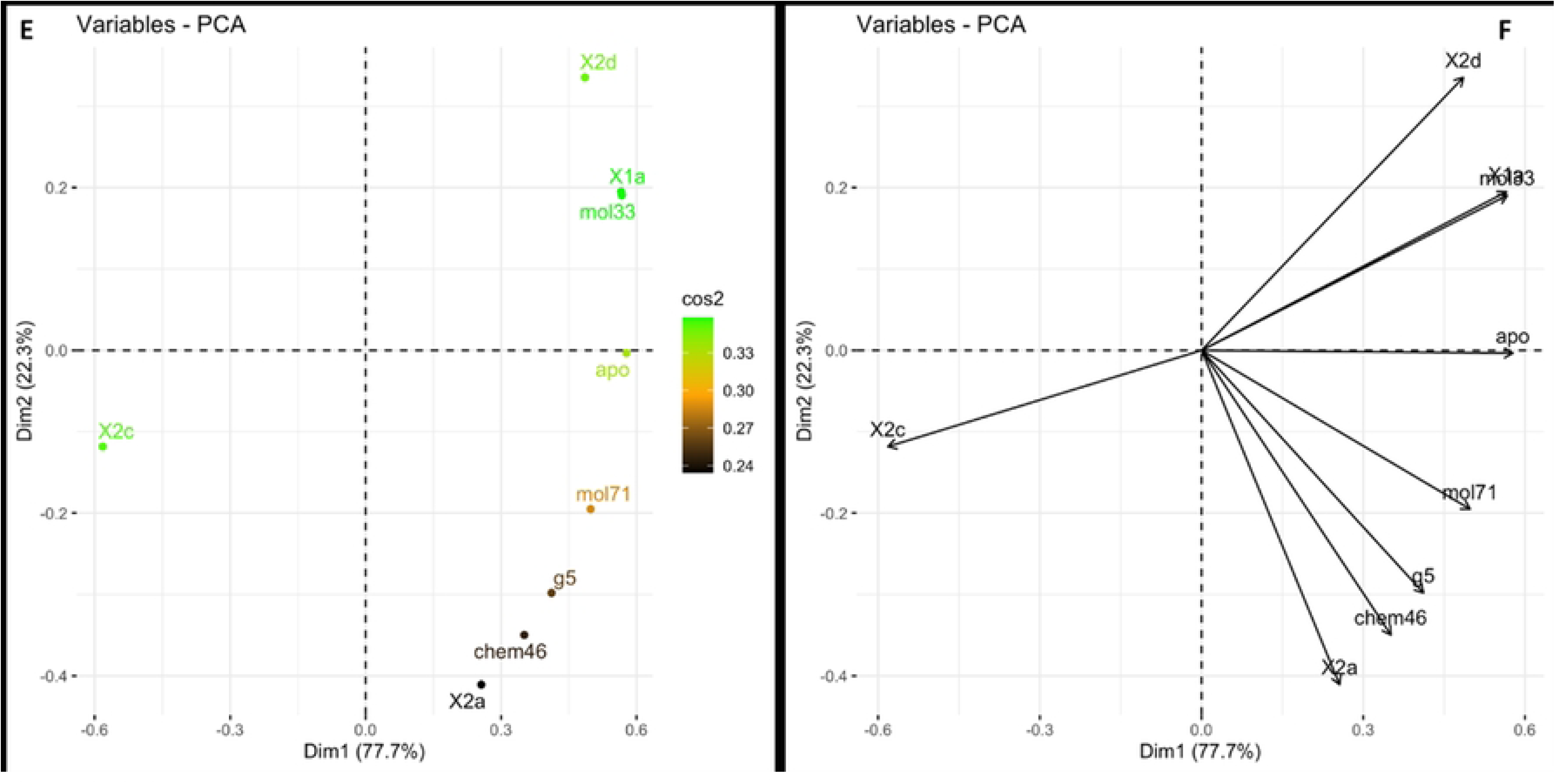
Statistical analysis of Cα residue fluctuations of residues in the USP18 domains during 100ns of simulation. A. Heatmap plot of RMSF of the IBB-1 residues. B. Heatmap plot of RMSF of the triad residues. C. Correlative PCA analysis of IBB-1 residues. D. Clustering PCA analysis of IBB-1 residues. E. Correlative PCA analysis of triad residues. F. Clustering PC analysis of triad residues.

By extracting residue fluctuations and measuring concerted changes in atom distances between residues in the domains during the NPT molecular dynamic simulations the induced-fit effect on critical residues of either the IBB-1 domain or the catalytic triad was monitored (Figs 6 and 7). The trajectories from the NPT simulations of the 8 ligand systems were compared to those of the unbound **apo** protein. PCA plots show that complexes with **2c** and **Mol33** cluster their IBB-1 domain residue fluctuations in the same quadrant (Fig 6 A, C and D). The plots of the triad domain show an anti-correlative clustering of **2c** with **Mol33** or the **apo** systems in contradiction to that observed with the IBB-1 domains (Fig 6 B, E and F). Interestingly ligand **2c** is having an impact on residues in both the IBB-1 domain and the catalytic triad, whereas **G5** IBB-1 fluctuations overlap the **apo** protein. This observation strengthens the case that the source of USP18 activity of **2c** and **G5** is likely due to impacts on the triad residues. It is likely that the position in the active pocket that **2c** occupies relative to **Mol33** and **G5** as well as the type of interactions that **2c** initiates in its binding lead to the observations of the differences in its impacts on the 2 domains we interrogated.

**Fig 7:**
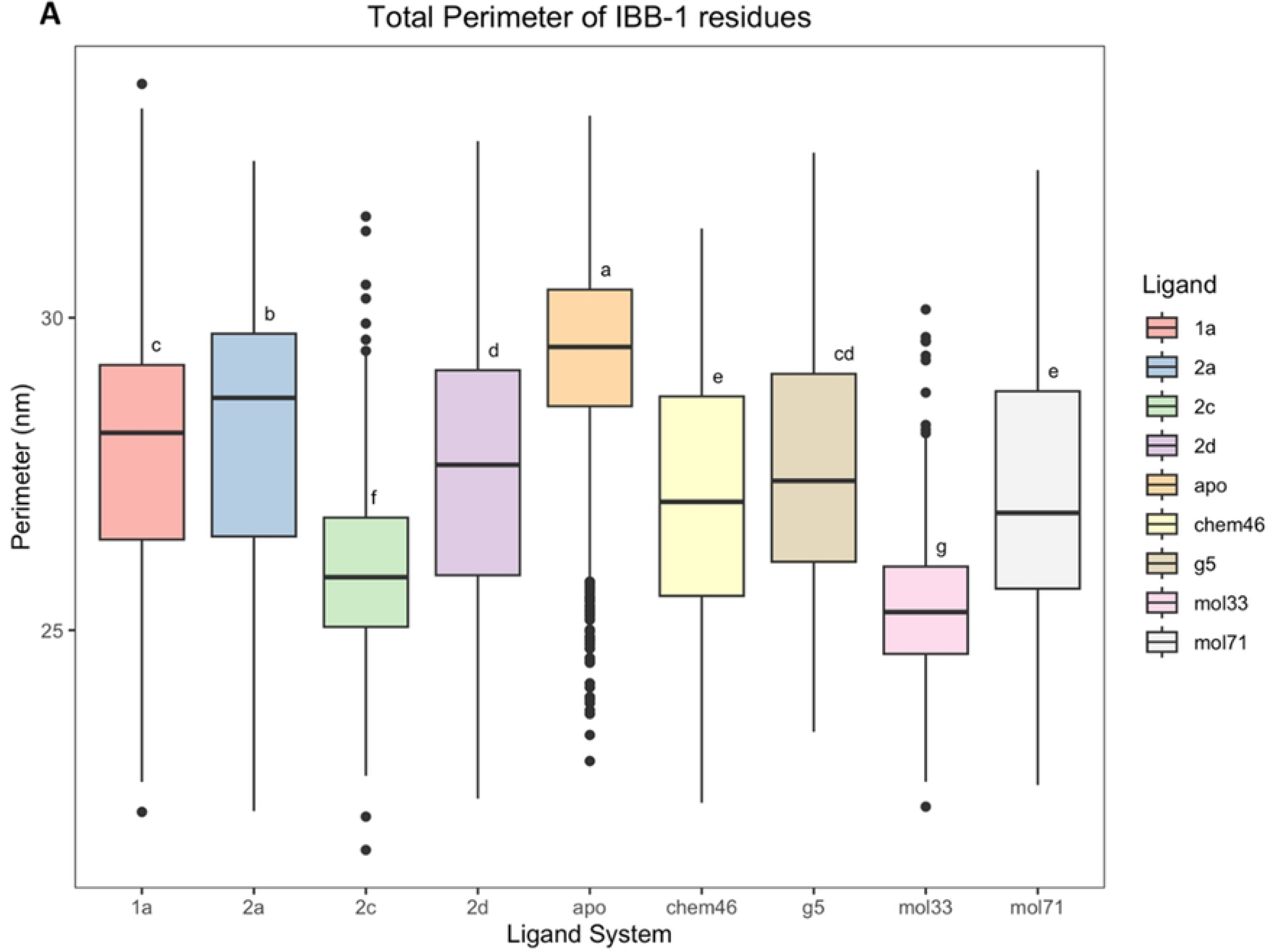
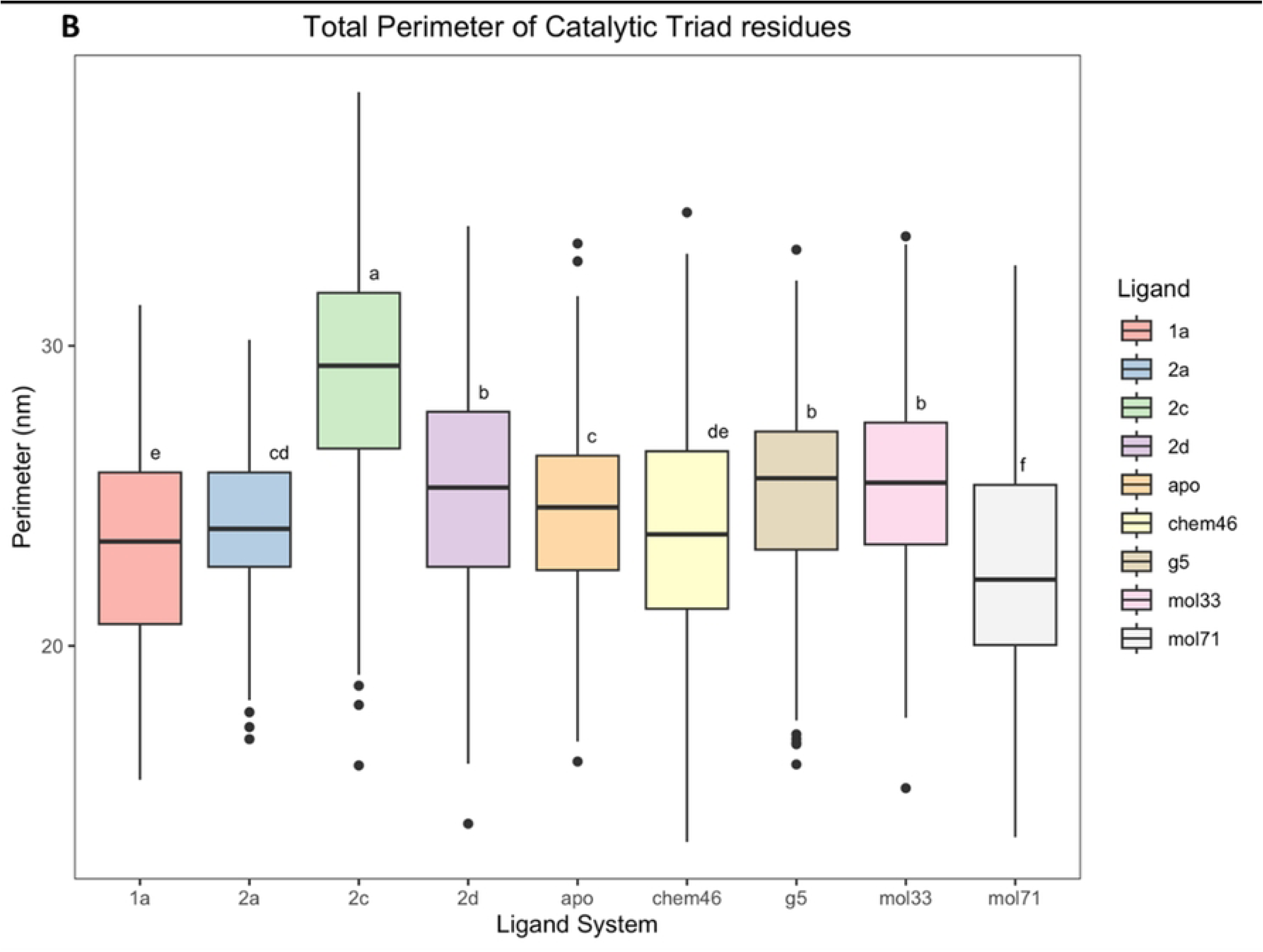
Total changes of atom distances within domain. A. Concerted IBB-1 domain atom distance changes. B. Concerted catalytic triad domain atom distance changes.

The distance changes between specific atoms of domain residues were collected as a single observable, total perimeter, allowing us to monitor the concerted distance change of the conformations at each time step. This analysis is analogous to measuring the changes in the gaps between the top and bottom bellows of an accordion over time. This observable allows us to deconvolute the relationships between the systems obtained from the multivariate PCAs. From the concerted total perimeter measurements, the domain residues of the systems show a correlation with ligand binding at the allosteric site (Fig 7). The IBB-1 residues show that conformations in ligand bound systems decrease the size of the total perimeter of this domain when compared to the unbound **apo** system (Fig 7 A). Ligand **2c** and **Mol33** that cluster in the same quadrant from their PCA (Fig 6 C and D), have the largest decrease in the total perimeter between these residues with the mean of **Mol33** (24.81 nm) lower than that of **2c** (26.25 nm) (Fig 7 A). The USP18 active ligand **2c** demonstrates a greater decrease in the mean of the total perimeter than **G5** (27.40 nm) consistent with the reported superior isopeptidase activity. These simulations put forward the role of allosteric inhibition in manipulating the IBB-1 domain from the induce-fit effect of ligand binding as the source of the isopeptidase activity. **Mol33**, which has a noticeable impact on the IBB-1 domain does appear to influence viral turnover to the same degree as **2c** from the anti-HIV-1 assays. The binding of all the small molecules has an impact on this IBB-1 domain and as such it is plausible that the anti-USP18 isopeptidase activity of **2c** and **G5** occurs due to impaired interactions with the ISGylated substrate. Binding affinity surface plasmon resonance studies, beyond the scope of this study, conducted in the presence of inhibitor could further elucidate whether the mechanism of impaired USP18 binding to ISGylated substrate is the source of the observed isopeptidase activity of **2c** and **G5**.

Close inspection of the triad residues suggests that the differences in the fluctuations of the **2c** system (Fig 6 B, E and F) occur due to the expansion of the catalytic triad (Fig 7 B). This opening results in a greater separation of the residues in the triad reflected in an increase in the RMSFs for the **2c** system not seen in the other ligand systems. Based on the total perimeter (Fig 7 B) **Mol33**, shared similarities with **2d** and **G5** (b), whereas when the magnitudes of the residue fluctuations were interrogated independent of direction and orientation through PCA, it overlapped with **1a** (Fig 6 E, F). Therefore, the magnitude of the triad contractions of the systems gives more resolution on the nature of the perturbation, than is shown by the PCA clustering.

Analysis of the simulations show that the induced-fit effect on the IBB-1 domain residues due to allosteric binding of all the ligands has a greater potential to explain the USP18 inhibition. Although the anti-HIV-1 assays did not equivocally confirm our hypothesis, it is possible that the low anti-HIV-1 activity we are observing is due to low anti-USP18 activity as is seen in **2c** and **G5**. We propose that in instances where USP18 deISGylation of ISGylated substrate viral proteins is inhibited successfully by small molecules, a decrease in the viability of critical viral proteins would be expected to result in the amplified detection of anti-HIV-1 activity from the assays. While the focus has been on the impact of the induced-fit effect on USP18’s isopeptidase activity, it is also possible that the pronounced induced-fit effect observed at **2c** at the allosteric site contributes to interferon signaling by reducing the IFNAR2 receptor binding. This would have the impact of influencing the USP18 function in an isopeptidase independent function. This interaction cannot be ruled out as residues on USP18 responsible for binding to STAT2 overlap with the residues of the catalytic triad domain that regulate isopeptidase activity (63). In this scenario however, if inhibitors such as **2c** cause a reduction in the binding of USP18 to IFNAR2 they would serve to reduce the non-enzymatic IFN dampening function of USP18. This reduction would allow the IFN cascade to proceed, as the phosphorylation of STAT1 and STAT2 would be prolonged resulting in the expression of antiviral chemokines and genes such as ISG15 as well as pro-viral USP18 (64). However, mutation studies using selective inhibition of USP18 isopeptidase activity have repeatedly led to enhanced viral resistance in mice due to increased ISGylation with minimal risk of severe side effects (65). Whether the isopeptidase inhibition is due to the allosteric induced-fit effect on the triad residues, the IBB-1 domain or is due to the reduction in IFN dampening is further complicated by the observation that it could be a combination of all three or two phenomenon. Elucidating the precise source of the elimination of viral proteins from these assays is beyond the scope of this study owing to the low activity of the USP18 inhibitors we have access to currently. It is possible that microarray studies of these phenomenon could confirm our suspicions and shed light on the mechanism of viral control due to allosteric binding (66).

### Free-energy profiling

Multi-trajectory simulations allowed free-energy profiling of the binding of the ligands at the allosteric site 2 of USP18. These profiles would assist in confirming whether patterns that explain the increased USP18 activity of **2c** over **G5** could be extrapolated to the other candidates based on their relationship with **G5** (Fig 9). It is also possible that the proposed mechanism of allosteric binding and the free-energy profiles of the small molecules could account for differences in the domain fluctuations (Fig 6) and the anti-HIV-1 activity observed (Fig 9 E).

Although **1a** (10.80 kcal mol^-1^) has a comparable PMF to that of **2c** (10.40 kcal mol^-1^) (Fig 8 D), the friction plot shows that **2c** dissociates from the protein later than **1a** (Fig 8 C). This later dissociation accounts for the increased induced-fit effects on the IBB-1 domain and the triad residues by **2c** (Fig 7). Although the PMF of **Mol33** (8.20 kcal mol^-1^) is less than **2c**, it has a comparable dissociation time. These multi-trajectory free-energy profiles show that **2c** has a comparable profile to that of **1a**, **2d** and **Mol33**. The superior isopeptidase activity of **2c** over **G5** can be attributed to combination of its superior binding affinity and the induced-fit effect of **2c** on the IBB-1 domain residues of USP18 which would be expected to translate into an improved anti-HIV-1 activity over **G5**. A combination of these traits is reflected across the set of compounds that we have evaluated in this study.

**Fig 8:**
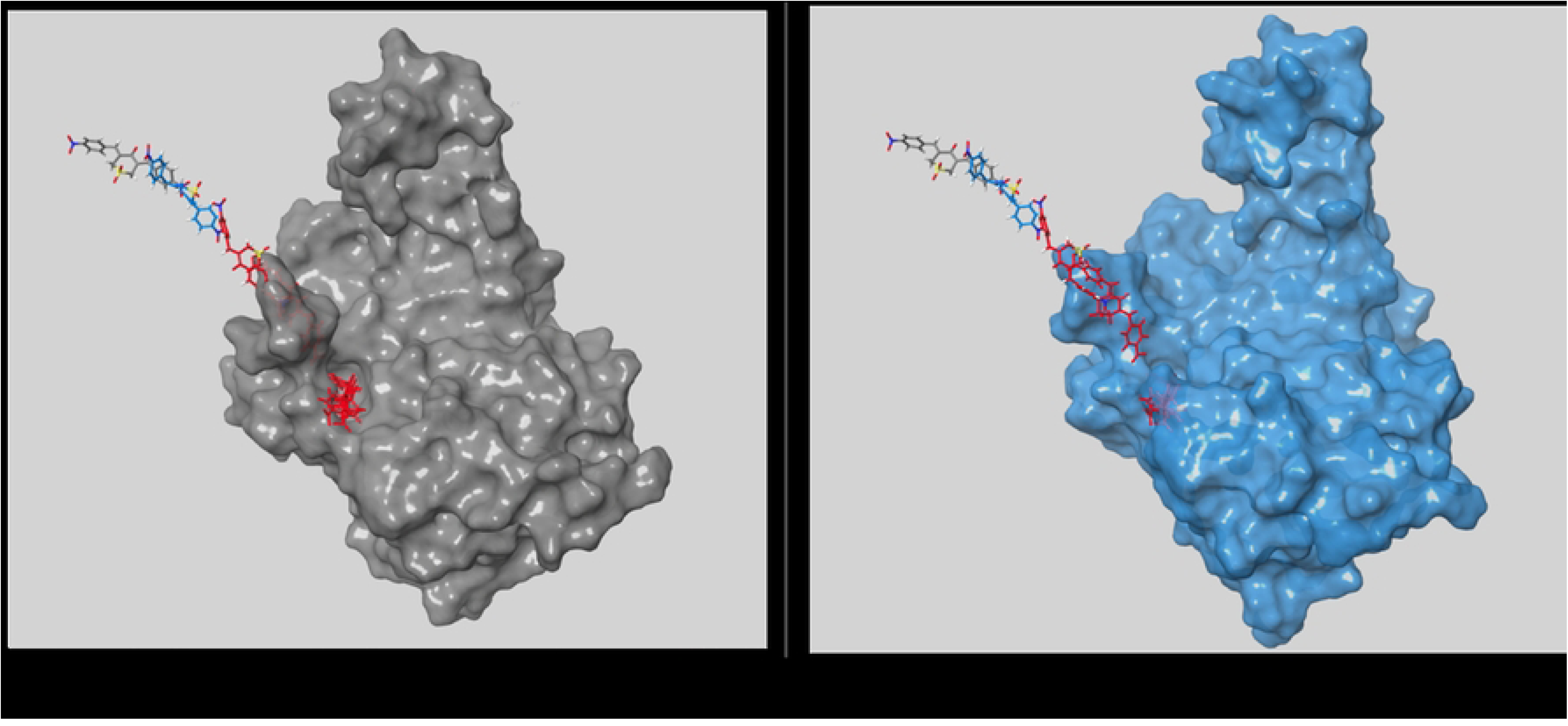
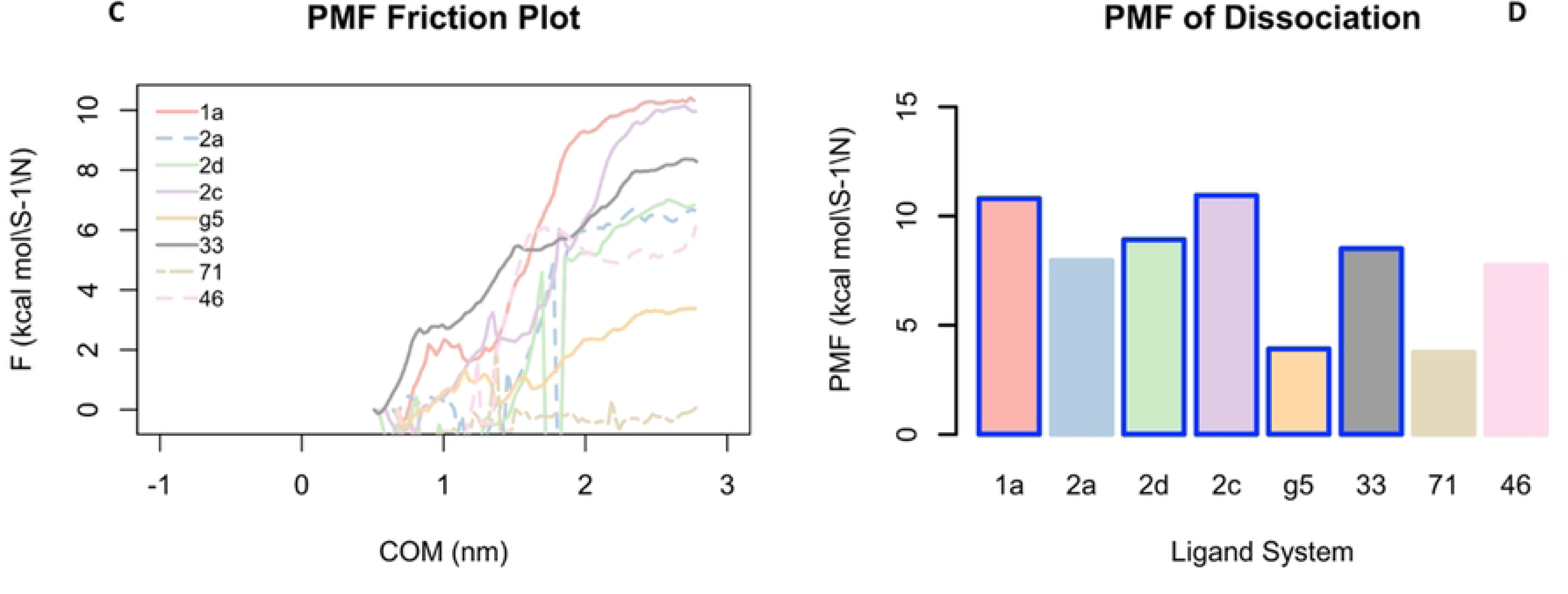
Free-energy profiling of ligand systems. A. Ligand G5 dissociation path with protein conformation at initial. B. Ligand G5 dissociation path with protein conformation after a few steps. C. PMF changes with COM distance. Solid lines are for systems obtained from 0.12 nm separation. D. Maximum PMF measured during dissociation for the 8 ligand systems. Boxes with borders show the systems obtained from 0.12 nm separation.

### Proof of concept for anti-viral activity with HIV-infected primary macrophages

The 5.2 million ZINC15 compounds that satisfied the criteria of MW between 325 and 425 g/mol, 3D representations obtained at pH 7.4, anodyne reactivity patterns, purchasable, and charged between +2 and -2 were screened against USP18 affinity using Glide XP with a grid box centered on site 2. The top 200 ligands were prioritized by ranking the compounds according to their Glide scores. For this study, 18 of the top 200 compounds that were immediately available for purchase from vendors at a satisfactory quality and located within ideal physicochemical space were purchased as the active ZINC15 set of compounds. Anti-HIV assays were performed on the 31 compounds (13 pyranone derivatives and the 18 USP18 active ZINC15 compounds) in human monocyte derived macrophages (hMDMs).

From the anti-HIV-1 assays, the bis-aryl pyranone compounds **G5**, **2c** and **2d** recorded activity of up to 60 % after 3 days post infection (dpi) and 70 % after 6 dpi for **2d** (Fig 9 B). This activity demonstrated the first proof of concept observation of the hypothesis, that anti-USP18 activity can lead to anti-HIV-1 activity. When the activity in the Alamar Blue assay was interrogated the **2d** activity coincided with a drop in cell viability whereas the activity of **2c** and **G5** was attributed to viral clearance (Fig 9 D). Inhibition of viral activity of 40 % by **G5** and 2c was more significant earlier in the infection cycle (3 dpi) and dropped to 20 % later in the infection cycle (6 dpi). Dose dependent plots after 3 dpi highlight how the cytotoxic effect of the pyranone derivatives (blue line) correlates with the anti-HIV activity (red line) observed (Fig 9 E). The ZINC filtered compounds on the other hand had modest inhibition activities of up to 20 % but eliminated the cytotoxicity of the pyranone derived compounds. The HIV screening assay demonstrates the utility of resolving genuine, non-cytotoxic, HIV inhibition. The promising activities serve as a proof of concept for the pursuit of anti-HIV small molecules targeted towards anti-USP18 activity. The impact of binding **Mol33** to the allosteric USP18 site on the fluctuations of key residues, and the concerted perturbations and expansions of functional domains, coupled with its binding free-energy profile, and non-cytotoxic anti-HIV inhibition underscore the need to explore the potential of accessing anti-HIV activity through attenuation of USP18 activity.

**Fig 9:**
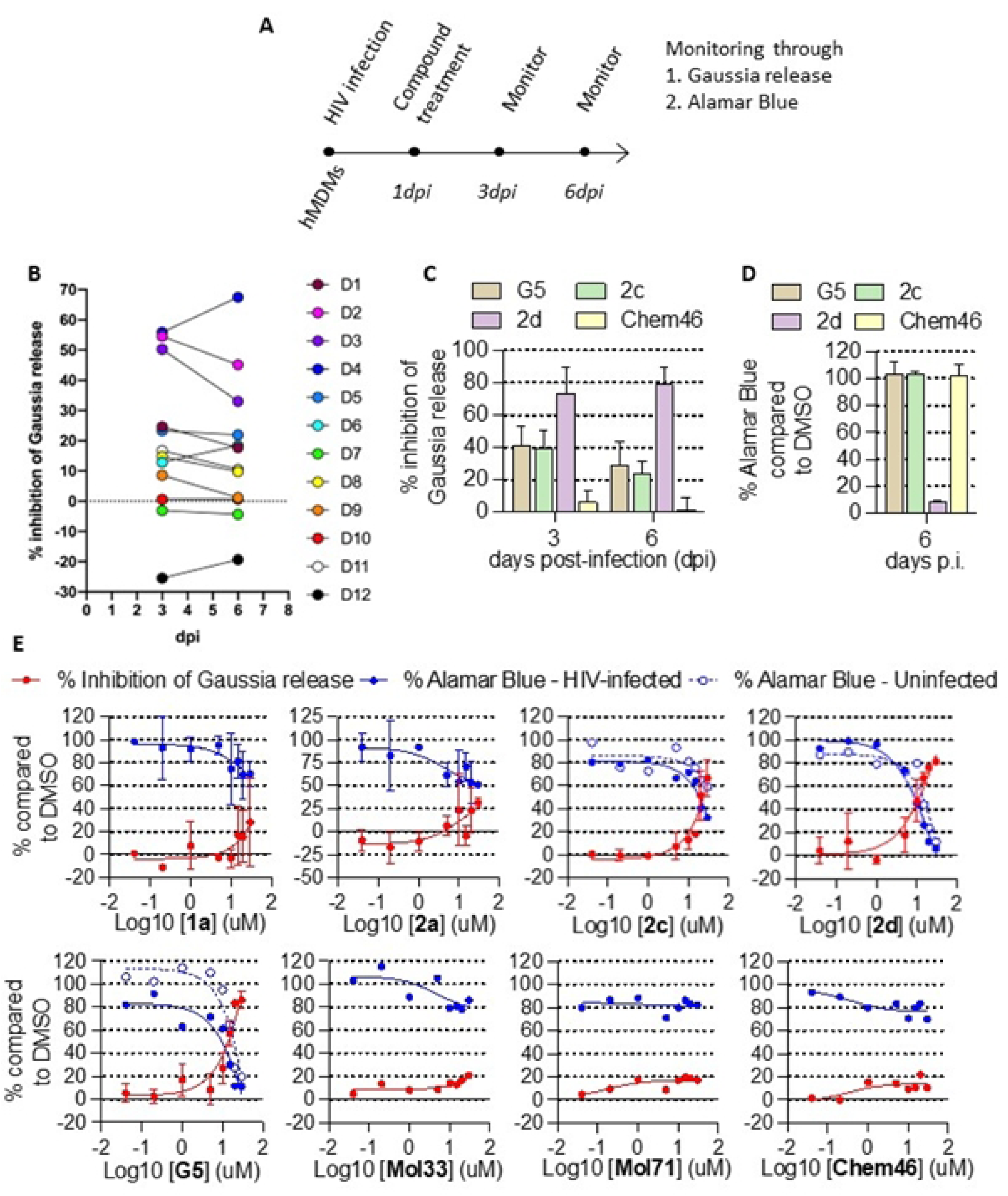
Compound screening for anti-HIV-1 activity in human monocyte-derived macrophages (hMDM). A. Timeline for compound screening B. Inhibition of HIV-1 viral production of 12 compounds D1-D12 measured by the inhibition of Gaussia release 3- or 6-days post infection (dpi). C-D. Inhibition of HIV-1 viral production measured by the inhibition of Gaussia release (C), and the cell viability compared to DMSO measured by the Alamar Blue Assay (D) of 4 compounds at 3 or 6 days post-infection (dpi) with 1 donor and 3 technical replicates. E. Dose dependent curves showing the inhibition of Gaussia release (in red) and Alamar Blue (in blue) for uninfected (circles) and HIV-infected macrophages (dots) recorded after 3 dpi and 2 days of bisaryl pyranones compounds G5, 2c, 2d, 1a and 2a; and best ZINC filtered compounds Chem46, Mol33 and Mol71. One to 2 donors are represented.

Although we have modest anti-HIV-1 activities, trends from molecular docking, free-energy profiles, and the influence of the induced-fit effect on the IBB-1 domain and the catalytic triad residues show that binding of small molecules at the allosteric site of USP18 impact its function and have the potential to direct the anti-HIV-1 activity. We postulate that the most dominant influence of anti-HIV-1 activity is the induced-fit perturbations on the USP18 IBB-1 domain residues arising from ligand binding at the allosteric site. We are not able to eliminate the contribution that large perturbations on the triad group due to expansion of this triad as this is beyond the scope of this study. The combined influence of these perturbations affects the activity of USP18 justifying the reported isopeptidase inhibition. We anticipate that inhibition in the early stages of infection enhance viral clearance by triggering ubiquitin-proteosome degradation of ISGylated critical viral proteins.

## Conclusions

We report the first attempt at acquiring anti-HIV activity from small molecules that target the IFN response by the perturbation of USP18, a key ISG factor in the interferon pathway. Bis-aryl pyranones with low micromolar USP18 activity were repurposed for anti-HIV activity. These antagonists were used to motivate for the hypothesis that fine-tuning the IFN response enhances natural control mechanisms which can lead to viral clearance. The pyranone derivatives had unfavourable metabolic and toxicity profiles as their scaffold possesses numerous structural alerts that render them unsuitable for medicinal development. Despite these drawbacks this series of compounds gives valuable insights into the role of USP18 antagonism as a promising strategy for novel anti-HIV therapies. A human USP18 model was acquired and metalated with Zn^2+^. This model was mapped for potential small molecule perturbation sites and an allosteric site that had ideal characteristics for small molecule binding was identified. The centre of this site overlapped with the centroid of residues **Ala299**, **Asp124**, **Gln122** and **Tyr348** on the human USP18 sequence from the Q9UMW8 homology model. Molecular dynamics studies highlighted the impact of ligand binding to this site. The analysis of reported active USP18 agonists that had anti-HIV-1 activity suggested that these inhibitors possess a reversible non-competitive inhibitory effect due to an induced-fit effect on domain residues that regulate its activity. Although the binding of active inhibitors at the allosteric site results in the expansion of the catalytic triad it is thought that the main driver for the isopeptidase inhibition is likely perturbation of the IBB-1 domain. The adoption of this conformation would reduce the turnover of cleaved ISG15 products within the cytosol resulting in an increase in ISG15 tagged proteins which was observed by Cersosimo (2015). It is our opinion that these ISG15-tagged proteins trigger the ubiquitin code resulting in the rapid degradation of the targeted viral proteins. Multi-trajectory simulations that allow the computation of the relative binding free-energy showed that the anti-HIV-1 activity of these anti-USP18 inhibitors was not attributable to a distinctively superior affinity to the USP18 allosteric pocket. The resulting anti-HIV-1 activity of these USP18 antagonists is thus due to a combination of induced-fit effects on the catalytic triad domain and the IBB-1 domain due to binding of small molecules at the allosteric site.

Novel anti-HIV scaffolds that possess potent anti-USP18 activity would need to be developed as the pyranone scaffolds possess undesirable toxicity profiles (Supplementary Information). This study suggests that a compound screening approach that considers the induce-fit effect is critical. This is a key determinant on anti-HIV-1 activity due to the impact of the catalytic functions of USP18 on the interferon signal cascade. Although inhibitors that mediate the type I IFN anti-viral response have been pursued in the past with limited success, this is the first study that has elucidated the specific role that USP18 inhibition and the ubiquitin-proteasome could play in this application. The allosteric inhibition of USP18 gives rise to an induced-fit effect with potential to influence both the enzymatic and non-enzymatic activities of USP18. This study shows that targeting the catalytic USP18 activity is a viable strategy to fine-tune the IFN response to enhance natural control mechanisms that lead to clearance of viral infection and a potential cure if targeted to reservoir cells.

## Acknowledgements

We acknowledge the Centre for High Performance Computing (CHPC), South Africa, for providing computational resources to this research project.

## Notes

### Competing Interest Statement

The authors have declared no competing interest.

## References

1. Netanya Utay AS, Douek DC. Interferons and HIV Infection: The Good, the Bad, and the Ugly. Pathogens and Immunity. 2016;1(1).

2. Veazey R, Pilch-Cooper H, Hope T, Alter G, Carias A, Sips M, et al. Prevention of SHIV transmission by topical IFN-β treatment. Mucosal immunology. 2016 Nov 1;9(6):1528–36.

3. Martins L, Szaniawski M, Williams E, Coiras M, Hanley T, Planelles V. HIV-1 Accessory Proteins Impart a Modest Interferon Response and Upregulate Cell Cycle-Related Genes in Macrophages. Pathogens (Basel, Switzerland). 2022 Feb 1;11(2).

4. Strebel K. HIV Accessory Proteins versus Host Restriction Factors. Current opinion in virology. 2013 Dec;3(6):692–9.

5. Su L. Pathogenic role of type I interferons in HIV-induced immune impairments in humanized mice. Current HIV/AIDS reports. 2019 Jun 15;16(3):224.

6. Ickler J, Francois S, Widera M, Santiago ML, Dittmer U, Sutter K. HIV infection does not alter interferon α/β receptor 2 expression on mucosal immune cells. PLOS ONE. 2020;15(1).

7. Xie X, Karakoese Z, Ablikim D, Ickler J, Schuhenn J, Zeng X, et al. IFNα subtype-specific susceptibility of HBV in the course of chronic infection. Frontiers in Immunology. 2022 Oct 14;0:6139.

8. Raman EP, Lakkaraju SK, Denny RA, MacKerell AD. Estimation of relative free energies of binding using pre-computed ensembles based on the single-step free energy perturbation and the site-identification by Ligand competitive saturation approaches. Journal of Computational Chemistry. 2017;

9. Li J, Koh JJ, Liu S, Lakshminarayanan R, Verma CS, Beuerman RW. Membrane active antimicrobial peptides: Translating mechanistic insights to design. Frontiers in Neuroscience. 2017.

10. Mishra A, Mulpuru V, Mishra N. Exploring the mechanism of action of podophyllotoxin derivatives through molecular docking, molecular dynamics simulation and MM/PBSA studies. 101080/0739110220222138549. 2022;

11. Basters A, Geurink PP, Röcker A, Witting KF, Tadayon R, Hess S, et al. Structural basis of the specificity of USP18 toward ISG15. Nature Structural and Molecular Biology. 2017;24(3).

12. Durfee L, Lyon N, Seo K, Huibregtse J. The ISG15 conjugation system broadly targets newly synthesized proteins: implications for the antiviral function of ISG15. Molecular cell. 2010 Jun 11;38(5):722–32.

13. Dzimianski J V., Scholte FEM, Bergeron É, Pegan SD. ISG15: It’s Complicated. Journal of Molecular Biology. 2019 Oct 4;431(21):4203–16.

14. Thery F, Eggermont D, Impens F. Proteomics Mapping of the ISGylation Landscape in Innate Immunity. Frontiers in Immunology. 2021 Aug 10;0:3089.

15. Kang JA, Jeon YJ. Emerging Roles of USP18: From Biology to Pathophysiology. International Journal of Molecular Sciences 2020, Vol 21, Page 6825. 2020 Sep 17;21(18):6825.

16. Liu Z, Bethunaickan R, Huang W, Lodhi U, Solano I, Madaio MP, et al. Interferon-α accelerates murine systemic lupus erythematosus in a T cell-dependent manner. Arthritis and Rheumatism. 2011 Jan;63(1):219–29.

17. Oon S, Wilson NJ, Wicks I. Targeted therapeutics in SLE: emerging strategies to modulate the interferon pathway. Clinical & Translational Immunology. 2016 May 1;5(5):e79.

18. Broering R, Zhang X, Kottilil S, Trippler M, Jiang M, Lu M, et al. The interferon stimulated gene 15 functions as a proviral factor for the hepatitis C virus and as a regulator of the IFN response. Gut. 2010 Aug 1;59(8):1111–9.

19. Luo H. Interplay between the virus and the ubiquitin–proteasome system: molecular mechanism of viral pathogenesis. Current Opinion in Virology. 2016 Apr 1;17:1.

20. Chen L, Li S, McGilvray I. The ISG15/USP18 ubiquitin-like pathway (ISGylation system) in Hepatitis C Virus infection and resistance to interferon therapy. The International Journal of Biochemistry & Cell Biology. 2011 Oct 1;43(10):1427–31.

21. Dagenais-Lussier X, Loucif H, Cadorel H, Blumberger J, Isnard S, Bego MG, et al. USP18 is a significant driver of memory CD4 T-cell reduced viability caused by type I IFN signaling during primary HIV-1 infection. PLOS Pathogens. 2019;15(10):e1008060.

22. Kim JH, Luo JK, Zhang DE. The Level of Hepatitis B Virus Replication Is Not Affected by Protein ISG15 Modification but Is Reduced by Inhibition of UBP43 (USP18) Expression. The Journal of Immunology. 2008 Nov 1;181(9):6467–72.

23. Cersosimo U, Sgorbissa A, Foti C, Drioli S, Angelica R, Tomasella A, et al. Synthesis, Characterization, and Optimization for in Vivo Delivery of a Nonselective Isopeptidase Inhibitor as New Antineoplastic Agent. Journal of Medicinal Chemistry. 2015 Feb 26;58(4):1691–704.

24. Bienert S, Waterhouse A, de Beer T, Tauriello G, Studer G, Bordoli L, et al. The SWISS-MODEL Repository-new features and functionality. Nucleic Acids Res [Internet]. 2017 Jan 1 [cited 2022 Sep 21];45(D1):D313–9. Available from: https://pubmed.ncbi.nlm.nih.gov/27899672/

25. Daina A, Michielin O, Zoete V. SwissADME: A free web tool to evaluate pharmacokinetics, drug-likeness and medicinal chemistry friendliness of small molecules. Scientific Reports. 2017;7.

26. Xiong G, Wu Z, Yi J, Fu L, Yang Z, Hsieh C, et al. ADMETlab 2.0: An integrated online platform for accurate and comprehensive predictions of ADMET properties. Nucleic Acids Research. 2021;49(W1).

27. Wishart DS, Tian S, Allen D, Oler E, Peters H, Lui VW, et al. BioTransformer 3.0 - a web server for accurately predicting metabolic transformation products. Nucleic Acids Research. 2022;50(W1).

28. Lin Y, Cheng C, Shih C, Hwang J, Yu C, Lu C. MIB: Metal Ion-Binding Site Prediction and Docking Server. J Chem Inf Model [Internet]. 2016 Dec 27 [cited 2022 Sep 21];56(12):2287–91. Available from: https://pubmed.ncbi.nlm.nih.gov/27976886/

29. Lu CH, Lin YF, Lin JJ, Yu CS. Prediction of Metal Ion–Binding Sites in Proteins Using the Fragment Transformation Method. PLoS One [Internet]. 2012 Jun 18 [cited 2022 Sep 21];7(6):e39252. Available from: https://journals.plos.org/plosone/article?id=10.1371/journal.pone.0039252

30. Halgren TA. Identifying and Characterizing Binding Sites and Assessing Druggability. J Chem Inf Model [Internet]. 2009 Feb 23 [cited 2022 Sep 21];49(2):377–89. Available from: https://pubs.acs.org/doi/full/10.1021/ci800324m

31. Schrödinger. LigPrep. New York, NY, USA; 2021.

32. Chemaxon. Chemaxon internal. 2023 [cited 2023 Mar 27]. MarvinSketch. Available from: https://marvinjs-demo.chemaxon.com/latest/demo.html

33. Cho AE, Guallar V, Berne BJ, Friesner R. Importance of accurate charges in molecular docking: Quantum Mechanical/Molecular Mechanical (QM/MM) approach. Journal of Computational Chemistry. 2005 Jul 15;26(9):915–31.

34. Friesner RA, Banks JL, Murphy RB, Halgren TA, Klicic JJ, Mainz DT, et al. Glide: A New Approach for Rapid, Accurate Docking and Scoring. 1. Method and Assessment of Docking Accuracy. Journal of Medicinal Chemistry. 2004;

35. Richard A. Friesner *, Robert B. Murphy †, Matthew P. Repasky †, Leah L. Frye ‡, Jeremy R. Greenwood †, Thomas A. Halgren †, et al. Extra Precision Glide: Docking and Scoring Incorporating a Model of Hydrophobic Enclosure for Protein−Ligand Complexes. Journal of Medicinal Chemistry. 2006 Oct;49(21):6177–96.

36. Irwin JJ, Shoichet BK. ZINC - A free database of commercially available compounds for virtual screening. Journal of Chemical Information and Modeling. 2005;

37. Sterling T, Irwin JJ. ZINC 15 - Ligand Discovery for Everyone. Journal of Chemical Information and Modeling. 2015;55(11).

38. Sherman W, Beard H, Farid R. Use of an induced fit receptor structure in virtual screening. Chemical biology & drug design. 2006;67(1):83–4.

39. Sherman W, Day T, Jacobson M, Friesner R, Farid R. Novel procedure for modeling ligand/receptor induced fit effects. Journal of medicinal chemistry. 2006 Jan 26;49(2):534–53.

40. Bowers KJ, Chow E, Xu H, Dror RO, Eastwood MP, Gregersen BA, et al. Scalable Algorithms for Molecular Dynamics Simulations on Commodity Clusters. In: Proceedings of the 2006 ACM/IEEE Conference on Supercomputing. New York, NY, USA: Association for Computing Machinery; 2006. p. 84–es. (SC ’06).

41. Banks JL, Beard HS, Cao Y, Cho AE, Damm W, Farid R, et al. Integrated Modeling Program, Applied Chemical Theory (IMPACT). Vol. 26, Journal of Computational Chemistry. 2005.

42. Lemkul JA, Bevan DR. Assessing the Stability of Alzheimer’s Amyloid Protofibrils Using Molecular Dynamics. Journal of Physical Chemistry B. 2010;114:1652–60.

43. Abraham MJ, Murtola T, Schulz R, Páll S, Smith JC, Hess B, et al. GROMACS: High performance molecular simulations through multi-level parallelism from laptops to supercomputers. SoftwareX. 2015 Sep 1;1–2:19–25.

44. Huang J, MacKerell A. CHARMM36 all-atom additive protein force field: validation based on comparison to NMR data. Journal of computational chemistry. 2013 Sep 30;34(25):2135–45.

45. Vanommeslaeghe K, Hatcher E, Acharya C, Kundu S, Zhong S, Shim J, et al. CHARMM general force field: A force field for drug-like molecules compatible with the CHARMM all-atom additive biological force fields. Journal of computational chemistry. 2010 Mar;31(4):671–90.

46. Vanommeslaeghe K, A. D. MacKerell Jr. Automation of the CHARMM General Force Field (CGenFF) I: Bond Perception and Atom Typing. Journal of Chemical Information and Modeling. 2012 Dec 21;52(12):3144–54.

47. Vanommeslaeghe K, Raman EP, A. D. MacKerell Jr. Automation of the CHARMM General Force Field (CGenFF) II: Assignment of Bonded Parameters and Partial Atomic Charges. Journal of Chemical Information and Modeling. 2012 Dec 21; 52(12):3155–68.

48. Hub JS, De Groot BL, Van Der Spoel D. G-whams-a free Weighted Histogram Analysis implementation including robust error and autocorrelation estimates. Journal of Chemical Theory and Computation. 2010;6(12).

49. Lipinski C a., Lombardo F, Dominy BW, Feeney PJ. Experimental and computational approaches to estimate solubility and permeability in drug discovery and development settings. Advanced Drug Delivery Reviews. 2012;64:4–17.

50. Ghose AK, Viswanadhan VN, Wendoloski JJ. A knowledge-based approach in designing combinatorial or medicinal chemistry libraries for drug discovery. 1. A qualitative and quantitative characterization of known drug databases. Journal of Combinatorial Chemistry. 1999;1(1).

51. Veber DF, Johnson SR, Cheng HY, Smith BR, Ward KW, Kopple KD. Molecular properties that influence the oral bioavailability of drug candidates. Journal of medicinal chemistry. 2002;45:2615–23.

52. Egan WJ, Merz KM, Baldwin JJ. Prediction of drug absorption using multivariate statistics. Journal of Medicinal Chemistry. 2000;43(21).

53. Muegge I, Heald SL, Brittelli D. Simple selection criteria for drug-like chemical matter. Journal of Medicinal Chemistry. 2001;44(12).

54. Geldenhuys WJ, Mohammad AS, Adkins CE, Lockman PR. Molecular determinants of blood-brain barrier permeation. Vol. 6, Therapeutic Delivery. 2015.

55. Roda E, Nion S, Bernocchi G, Coccini T. Blood-brain barrier (BBB) toxicity and permeability assessment after L-(4-10Boronophenyl)alanine, a conventional B-containing drug for boron neutron capture therapy, using an in vitro BBB model. Brain Research. 2014;1583(1).

56. Murakami T, Takano M. Intestinal efflux transporters and drug absorption Vol. 4, Expert Opinion on Drug Metabolism and Toxicology 2008.

57. Zhao M, Ma J, Li M, Zhang Y, Jiang B, Zhao X, et al. Cytochrome p450 enzymes and drug metabolism in humans. Vol. 22, International Journal of Molecular Sciences. 2021.

58. Brenk R, Schipani A, James D, Krasowski A, Gilbert IH, Frearson J, et al. Lessons learnt from assembling screening libraries for drug discovery for neglected diseases. ChemMedChem. 2008;3(3).

59. Magalhães PR, Reis PBPS, Vila-Viçosa D, Machuqueiro M, Victor BL. Identification of Pan-Assay INterference compoundS (PAINS) Using an MD-Based Protocol. In: Methods in Molecular Biology. 2021.

60. Lamothe SM, Guo J, Li W, Yang T, Zhang S. The Human Ether-a-go-go-related Gene (hERG) potassium channel represents an unusual target for protease-mediated damage. Journal of Biological Chemistry. 2016;291(39).

61. Nepali K, Lee HY, Liou JP. Nitro-Group-Containing Drugs. Vol. 62, Journal of Medicinal Chemistry. 2019.

62. Patterson S, Fairlamb AH. Current and Future Prospects of Nitro-compounds as Drugs for Trypanosomiasis and Leishmaniasis. Current Medicinal Chemistry. 2018;26(23).

